# Age-maintained human neurons demonstrate a developmental loss of intrinsic neurite growth ability

**DOI:** 10.1101/2023.05.23.541995

**Authors:** Bo P. Lear, Elizabeth A.N. Thompson, Kendra Rodriguez, Zachary P. Arndt, Saniya Khullar, Payton C. Klosa, Ryan J. Lu, Christopher S. Morrow, Ryan Risgaard, Ella R. Peterson, Brian B. Teefy, Anita Bhattacharyya, Andre M.M. Sousa, Daifeng Wang, Bérénice A. Benayoun, Darcie L. Moore

**Author notes:** Corresponding author, Darcie L. Moore. denotes equal contributions (Bo P. Lear, Elizabeth A.N. Thompson).

## Abstract

Injury to adult mammalian central nervous system (CNS) axons results in limited regeneration. Rodent studies have revealed a developmental switch in CNS axon regenerative ability, yet whether this is conserved in humans is unknown. Using human fibroblasts from 8 gestational-weeks to 72 years-old, we performed direct reprogramming to transdifferentiate fibroblasts into induced neurons (Fib-iNs), avoiding pluripotency which restores cells to an embryonic state. We found that early gestational Fib-iNs grew longer neurites than all other ages, mirroring the developmental switch in regenerative ability in rodents. RNA-sequencing and screening revealed ARID1A as a developmentally-regulated modifier of neurite growth in human neurons. These data suggest that age-specific epigenetic changes may drive the intrinsic loss of neurite growth ability in human CNS neurons during development.

**One-Sentence Summary:** Directly-reprogrammed human neurons demonstrate a developmental decrease in neurite growth ability.

## Main text

Human spinal cord injury (SCI) leads to life-long disability with high medical, psychological, and social costs (*1*). This injury damages long axonal projections originating from neurons in the brain, which extend through the spinal cord to communicate with the periphery. These central nervous system (CNS) axons fail to regenerate due to both inhibitory proteins in the injured environment, and by a lack of intrinsic axon growth ability in the neurons themselves (*2*). Interestingly, rodent studies have revealed a link between age and axon growth and regeneration, such that embryonic CNS neurons can extend long axons, while postnatal and adult CNS neurons cannot (*3–7*). Further, many developmentally-regulated genes, when modulated, can drive axon growth (*3, 8–13*). However, it is not known whether human neurons also have a developmental switch in CNS axon regenerative ability.

Here, we use an *in vitro*, age-retained, human neuronal model system to identify novel, developmentally-regulated, intrinsic factors which regulate neurite growth. To maintain the age of the originating fibroblasts, we utilized a validated direct reprogramming protocol which transdifferentiates human fibroblasts of different ages directly into induced neurons (Fib-iNs), avoiding pluripotency, which restores cells to an embryonic age (*14, 15*). In adult cells, the use of direct reprogramming maintains the age of the original cell, and has been shown in many different studies to reveal age-associated disease phenotypes (*14–20*), however whether prenatal age is similarly maintained with direct reprogramming is not clear. Additionally, direct reprogramming produces neurons which are environmentally naïve, allowing us to determine the role of age on the intrinsic neurite growth ability of human neurons.

Thus, to address these questions, we transdifferentiated human neurons from 11 fibroblast lines ranging from 8 gestational-weeks (GW) through 72 years-old (YO; Table 1) by overexpressing the transcription factors achaete-scute family bHLH transcription factor 1 (ASCL1) and neurogenin 2 (NEUROG2), and culturing cells in the presence of small molecules for 4 weeks (Fig. 1A-B, S1A-B) (*14, 15*). To first determine the types of neurons being made and the maintenance of age in Fib-iNs at the transcriptional level, we collected all ages of Fib-iNs for RNA-sequencing (RNA-seq). To obtain a pure neuronal population, we transduced Fib-iNs with a lentivirus expressing mScarlet from a hSynapsin promoter and used fluorescence activated cell sorting (FACS) to collect mScarlet+ neurons (Fig. S1C-D) to perform low cell number RNA-seq on all ages of Fib-iNs (Table 1). Samples were corrected for the 5 first genotype principal components (PCs), as determined from RNA-seq data (Fig. S1E), and examined for separation using multi-dimensional scaling (MDS), demonstrating that age-specific replicates clustered similarly (Fig. 1C, S1F).

**Figure 1:**
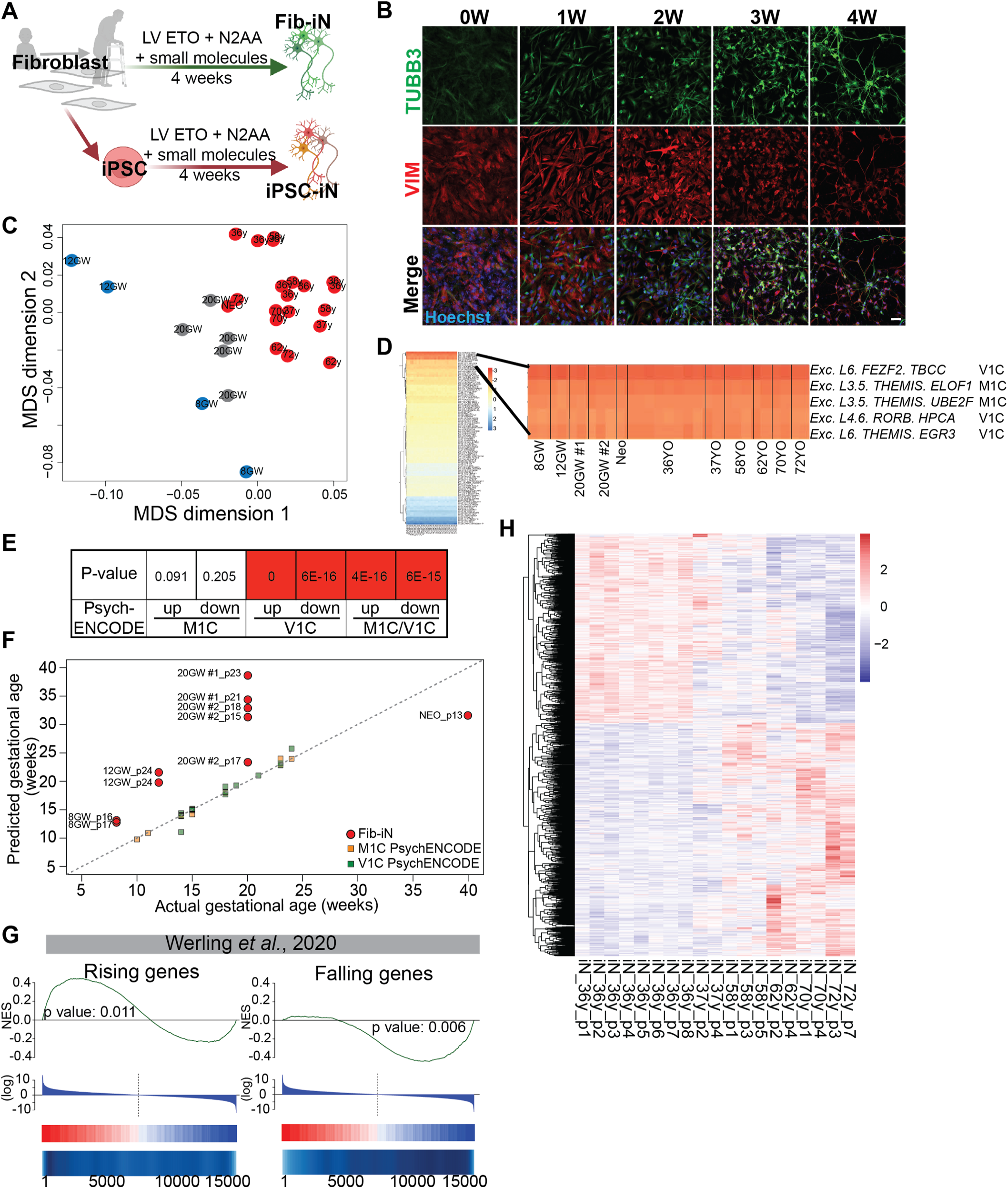
Fib-iNs transdifferentiated from fetal through adult human fibroblasts preserve age. **A-B**) 8GW to 72YO human fibroblasts were reprogrammed to Fib-iNs by overexpressing NEUROG2 and ASCL1 (N2AA) in a doxycycline-dependent manner (LV ETO), followed by 4 weeks of transdifferentiation in induction media. As an age-reset control, 5 of the original fibroblast ages were reprogrammed to iPSCs, then transdifferentiated using the same overexpression paradigm to generate iPSC-iNs. **B**) 36YO fibroblasts were immunostained over the 4 weeks of transdifferentiation to Fib-iNs for beta-III tubulin (TUBB3; neurons), vimentin (VIM; fibroblasts), and Hoechst (nuclei). Scale bar = 100 μm. **C**) Multi-dimensional scaling (MDS) of RNA-seq data for all Fib-iN samples. **D**) Spearman rank correlation (Rho) of Fib-iN RNA-seq to neuronal expression profiles from the Allen Brain Map single cell human data of various brain regions (*21, 22*), revealing that all ages of Fib-iNs are most similar to excitatory neurons (Exc.) of primary motor cortex (M1C) and primary visual cortex (V1C). **E**) Comparison of Fib-iN DEGs enriched in long-growing (8GW-12GW) vs. short-growing (NEO-72YO) against genes up and downregulated in human “M1C only”, “V1C only”, and “M1C and V1C” cortical development from PsychENCODE (*32*) with associated p-values. FDR <0.001 (V1C and V1C/M1C), FDR<0.013 (M1C UP), FDR<0.22 (M1C DOWN). **F**) Using machine-learning, RNA-seq from PsychENCODE’s human developmental M1C and V1C brain regions was used to train a model to predict the age of Fib-iN samples from RNA-seq profiles (*32*), revealing increasing developmental age from gestational age Fib-iNs, irrespective of passage number (p). **G**) Heatmap of significant age-related transcriptional changes in adult Fib-iNs (36YO-72YO; 2,431 genes) with an FDR<5%. H) Gene set enrichment analysis using genesets from human dorsolateral prefrontal cortex samples (*33*) showing genes rising during development (left) and genes falling in development (right) are enriched in short-growing (positive values) and long-growing Fib-iNs (negative values), respectively.

To confirm the neuron subtype specificity in all samples, we immunostained Fib-iNs for beta-III tubulin (TUBB3) together with either vesicular glutamate transporter 1 (VGLUT1; excitatory neurons) or gamma-aminobutyric acid (GABA; inhibitory neurons), and found that the neurons produced were 70.3% +/-6.7% VGLUT1+ and 16.5% +/-2.4% GABA+, similar to previously published reports (*14, 15*) (Fig. S1G-H), demonstrating that the neurons are primarily glutamatergic. To gain more detailed knowledge about the type of neurons being produced, we compared our RNA-seq data to the Allen Brain Map single cell RNA-seq data from adult human cortex (*21, 22*). We again confirmed that all ages of Fib-iNs made excitatory neurons, and the same neuron identity was conserved across the 11 lines (Fig. 1D, Table 1). Specifically, the Fib-iN transcriptional profiles were most similar to excitatory cortical neurons, with the topmost hit being excitatory neurons from layer 6, with FEZ family zinc finger 2 (*FEZF2*) and Tubulin Folding Cofactor C (*TBCC*) as their primary subclass markers and cluster-specific markers, respectively (Fig. 1D). This expression signature was most similar to cells from the primary visual cortex (V1C), middle temporal gyrus (MTG), and cingulate gyrus (CgG). We also identified strong similarities between Fib-iNs to cells from layers 3 and 5 (identifiers: *THEMIS* (subclass), *ELOF1* (cluster-specific) or *UBE2F* (cluster-specific)), whose expression signature was most similar to neurons in the M1 lower limb region of primary motor cortex and M1 upper limb region of primary motor cortex (Fig. 1D). Thus, these analyses confirm that human Fib-iNs are not only excitatory, but resemble human cortical neurons expressing *FEZF2*, a gene required for corticospinal motor neuron identity in rodents (*23–25*).

Use of this model system not only delivers the benefit of producing human neurons, but direct reprogramming also maintains the cellular age of the original cell by bypassing pluripotency during the transdifferentiation (*14, 15, 26*). This protocol has primarily been used to produce adult neurons from adult fibroblasts for aging studies (*27*), however there is limited knowledge of the ability to maintain prenatal age during direct reprogramming. To confirm general maintenance of cellular age in the Fib-iNs produced from our fibroblast lines of gestational through adult ages, we first immunostained fibroblasts and Fib-iNs for the nuclear envelope marker LaminB1, a protein whose levels decrease with aging (*16, 28–31*). We found a significant decrease in Lamin B1 with increasing age in both fibroblasts and Fib-iNs, confirming maintenance of cellular age with direct reprogramming in our fibroblast lines (Fig. S1I-J). To confirm maintenance of prenatal ages in Fib-iNs, we used the PsychENCODE region-specific, bulk tissue, RNA-seq data of prenatal and postnatal human brain to compare to gestational age Fib-iNs (8GW, 12GW, 20GW#1, 20GW#2, and Neonatal) (*32*). First, we compared genes significantly up- or downregulated with age in Fib-iNs to genes up- or downregulated in human V1C or M1C, and found genes up- and downregulated in V1C were significantly similar to those in Fib-iNs (Fig. 1E). Further, comparison of genes that were similarly regulated between the two areas (M1C and V1C) to our Fib-iNs revealed a significant similarity between the human tissue data and purified prenatal Fib-iNs (Fig. 1E). Second, to further confirm the maintenance of prenatal ages using this direct reprogramming approach, we trained a machine-learning model using bulk gene expression studies from dissected human M1C and V1C, ages 10GW to 24GW (*32*), to predict developmental age based on gene expression patterns. Using this model we were able to predict the age of each of our Fib-iN samples from 8GW to neonate, and found, despite comparing purified cultured excitatory neurons (Fib-iNs) to bulk tissue made up of many cell types (PsychENCODE) (*32*), 8GW to neonatal directly reprogrammed Fib-iNs had increasing predicted age, though with a slight upwards shift of predicted age in the Fib-iNs (Fig. 1F).

Recent studies on gene expression changes during human brain development revealed that the largest change in gene expression occurs during the late-fetal transition, a period beginning around 21GW (*32–34*). The genes whose expression increases during this late-fetal transition exhibited increased expression levels in postnatal to aged Fib-iNs (Neo-72YO, p=0.011), whereas those genes whose expression decreases during the late-fetal transition demonstrated decreased expression in early fetal Fib-iNs (8GW-12GW, p=0.006; Fig. 1G). Importantly, these data taken together suggest that prenatal age can be maintained with direct reprogramming, and that gestational Fib-iNs may be used to study neuronal function associated with intrinsic changes occurring during development.

Use of direct reprogramming also maintained transcriptional age in adult Fib-iNs. We found that adult Fib-iNs demonstrated changes in gene expression associated with adult aging, as expected, with 2,431 genes significantly changed (FDR<5%; Fig. 1H). Additionally, we found an increase in transposable element (TE) expression with age (Fig. S1K), confirming previous reports that TEs levels increase with age (*35*). Thus, our RNA-seq results support maintenance of age in Fib-iNs at all ages, both during gestation and aging.

Rodent CNS neurons undergo a developmental loss of axon growth ability around the time of birth (*3, 4, 7, 11, 36, 37*). Thus, using this validated human, age-specific model system, we asked if there were differences in intrinsic neurite growth ability among the 11 ages of Fib-iNs (TUBB3+/GABA-/VIM-; Fig. S2A-B). Neurons were dissociated after 4 weeks of transdifferentiation and sorted onto a gelatin substrate to remove remaining fibroblasts and enrich for neurons (Fib-iNs), before plating at low density (100 cells/mm^2^) on a laminin-coated substrate (Fig. S2C) for 2 days in an axon supportive media (ASM; S2D-H). Interestingly, we found that 8GW and 12GW human neurons grew the longest neurites of all the ages. By 20GW and older, neurons grew only short neurites, demonstrating a developmental decline in neurite growth ability in human neurons (Fig. 2A-B). Interestingly, when comparing neurodevelopmental benchmarks between mouse and human, mouse birth (E21/P0) is equivalent to approximately human 21GW (*38*), the timing of the late-fetal transition (*33*), suggesting the conservation of a developmental loss of intrinsic neurite growth ability in our environmentally naïve human neurons (Fig. S2I). Previously, it has been suggested that during neurodevelopment there is a transcriptional shift from elongation to synaptogenesis and branching which limits axon growth ability in rodents postnatally (*39, 40*). Thus, we measured neurite branching in all ages of Fib-iNs and found that total branching was increased in adult Fib-iNs (Fig. 2C), but this change did not coincide specifically with the time when intrinsic neurite growth ability decreased, nor was there a relationship between the number of branches on the longest neurite and Fib-iN age (Fig. S2J). Thus, we found that human Fib-iNs demonstrate a developmental loss of intrinsic neurite growth ability even in an *in vitro*, environmentally naïve system (Fig. 2A-B).

**Figure 2:**
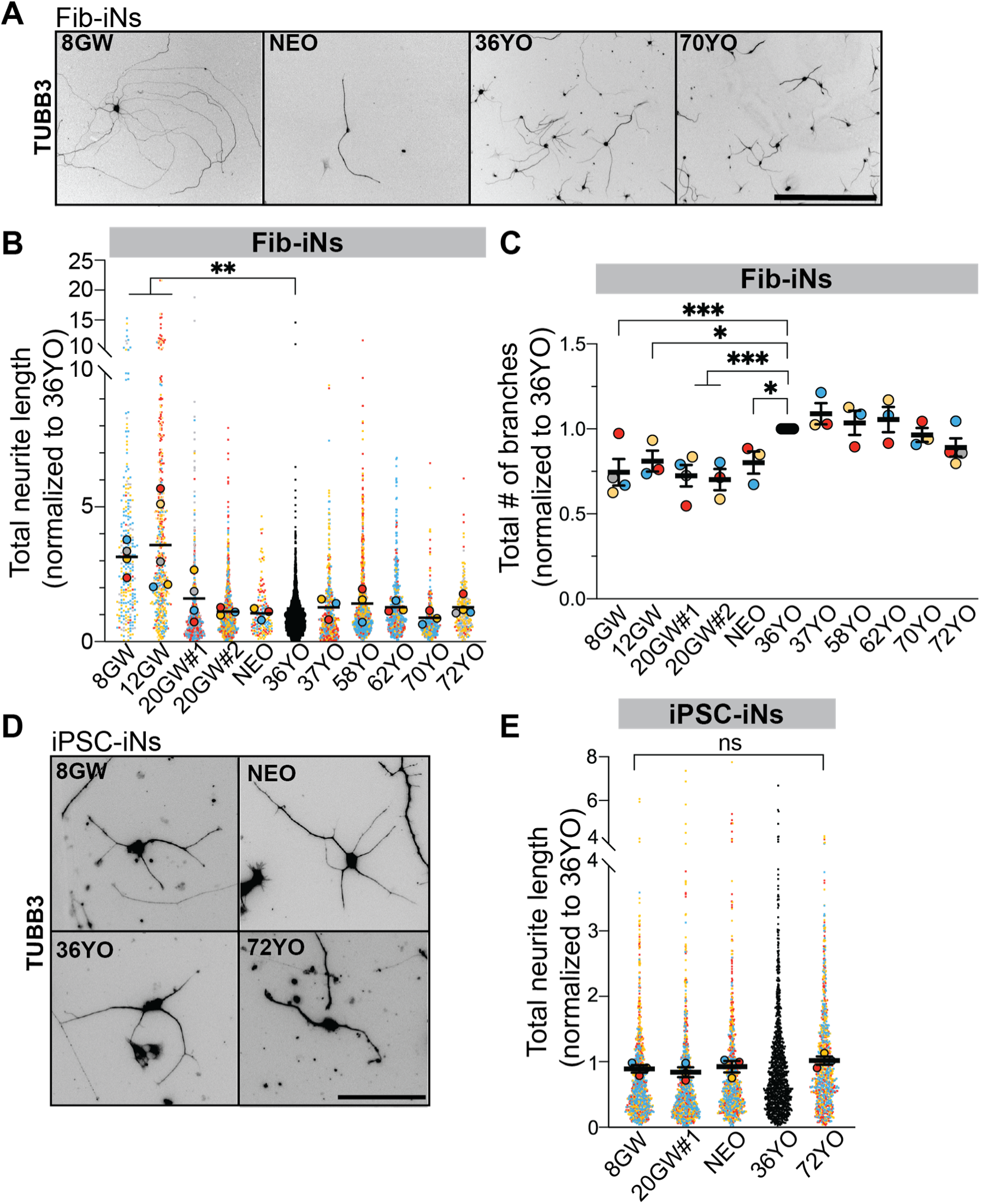
Conservation of a developmental loss of intrinsic neurite outgrowth in human neurons. **A-B**) Fib-iNs of all ages were trypsinized, and plated for 2 days in ASM before fixation, and immunostaining for TUBB3. Neurons were imaged and traced for total neurite length. A superplot shows total neurite length of all Fib-iN ages normalized to their within-batch 36YO sample. One-way ANOVA with post-hoc Tukey’s test on averages of replicates compared to 36YO; Mean ± SEM. **C**) The number of total branches per neuron were averaged from each age, and normalized to the 36YO sample of each experiment. One-way ANOVA with post-hoc Tukey’s test on averages of replicates compared to 36YO; Mean ± SEM. **D-E**) iPSC-iNs were trypsinized, and plated for 2 days in ASM before fixation, and immunostaining for TUBB3. A superplot shows total neurite length of all iPSC-iN ages normalized to their within-batch 36YO sample. One-way ANOVA with post-hoc Tukey’s test on averages of replicates; Mean ± SEM. For all experiments, N ≥ 3 different inductions, n ≥ 50 cells per induction. Scale bar (A) 500 μm, (E) 100 μm. ns = not significant, *p<0.05 **p<0.01 ***p<0.001.

To confirm that this neurite outgrowth phenotype was specific to the age of the Fib-iNs, we took 5 of the human fibroblast lines (Table 1), and reprogrammed them to iPSCs (Fig. 1A), thus resetting cellular age (Fig. S2K-L) as has been previously shown by others (*14–16, 41*). We directly reprogrammed these iPSCs to iNs (*14*) (iPSC-iNs; Fig. 1A, S2M-N), and found there was no longer any difference in the intrinsic neurite growth ability (Fig. 2D-E) of neurons between ages, demonstrating that the developmental decline in neurite growth ability in Fib-iNs is specific to the preservation of age.

Developmentally-regulated genes identified in rodents have been targeted to reveal age-dependent drivers of CNS axon growth and regeneration (*3, 11, 13, 42-44*). Thus, we asked which genes were significantly changed between “long-growing” Fib-iNs (8-12GW) versus “short-growing” Fib-iNs (Neo-72YO; Fig. 3A). Due to the neurite growth variability between the 20GW biological replicates and the variability in their “predicted” ages (Fig. 1G, 2B), we grouped both 20GW samples into a “variable condition,” and did not include them in this direct comparison. We identified 3,465 differentially expressed genes (DEGs) between long-growing and short-growing human Fib-iNs at FDR<1% (Fig. 3A, S3A, Table 2), indicating large transcriptional differences in the neurons with different growth phenotypes. Interestingly, using Gene Ontology enrichment analysis, we found enrichments for terms such as translational initiation, ribosomes, and cytoskeleton in long-growing Fib-iNs, whereas tissue-specific immune responses were enriched in short-growing Fib-iNs (Fig. S3A). We next asked how transcripts enriched in long-growing and short-growing Fib-iNs compare to known axon growth and regeneration datasets in rodent studies. Using gene sets from developmentally-regulated rodent CNS neuron studies, we compared the similarity of the transcriptional regulation of these genes to long- and short-growing Fib-iNs, revealing that human Fib-iNs behaved similarly to multiple developmentally-regulated gene sets in rodent retinal ganglion cells, cortical, and callosal neurons (Fig. 3B). However, a similar analysis using rodent axon regeneration datasets revealed less conservation (Fig. 3C). Finally, we curated a list of genes identified in other species to be activators or inhibitors of axon growth and regeneration (though not all developmentally-regulated) and assessed their expression levels across age in Fib-iNs (*33, 42, 43, 45-49*). Interestingly, we found only a subset of these genes tended to be differentially regulated with age in human Fib-iNs (Fig. S3B). These comparisons suggest that there are largely conserved developmental programs between species (Fig. 3B, S3B), however, regeneration programs have less agreement between rodent neurons and Fib-iNs (Fig. 3C, S3B). It is unclear whether this is due to the lack of orthologs between species for comparison, aspects of the Fib-iN protocol itself, or differences between humans and rodents.

**Figure 3:**
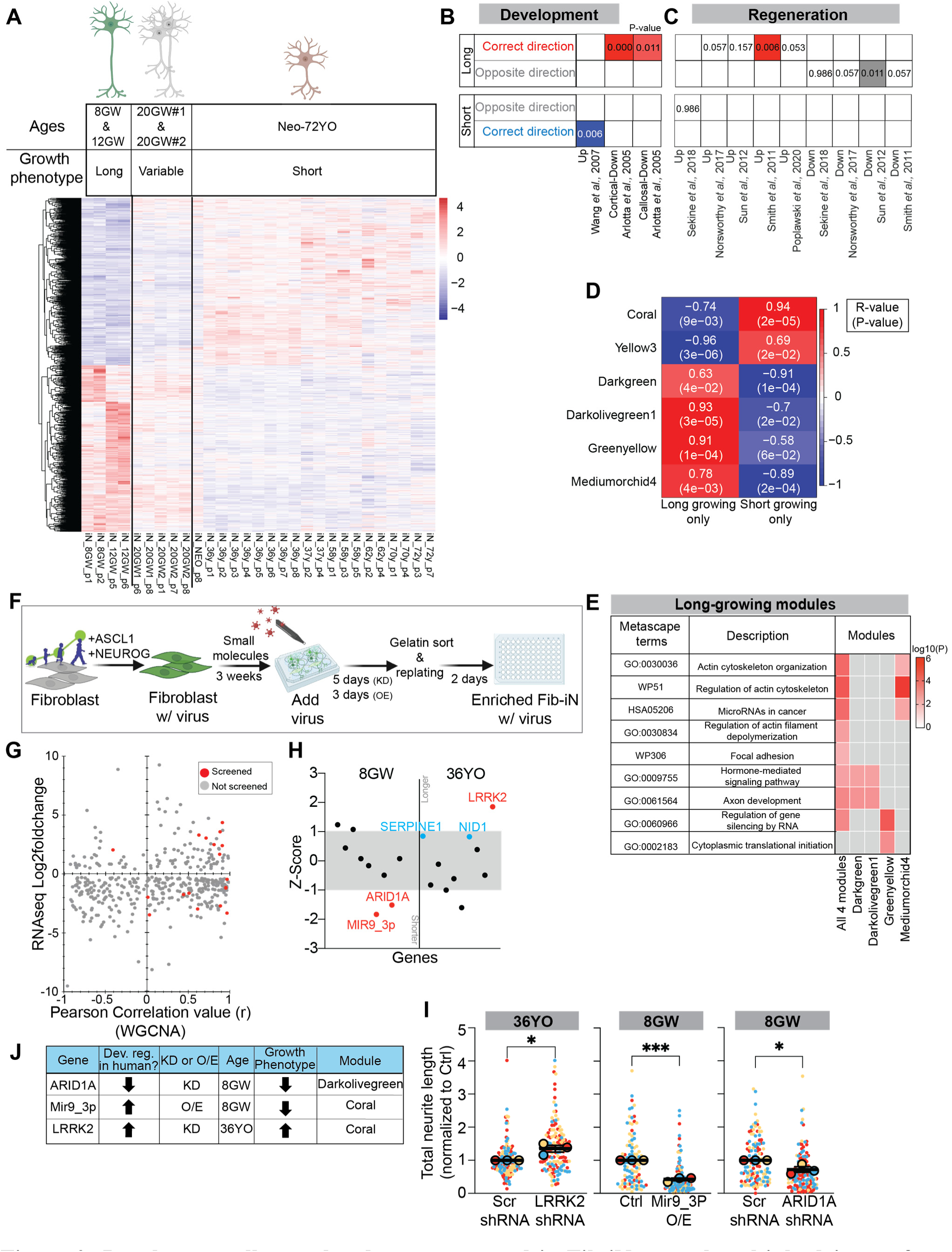
Developmentally-regulated genes screened in Fib-iNs reveal multiple drivers of human neurite growth. **A**) RNA-seq of 11 ages of Fib-iNs revealed 3,465 differentially expressed genes (DEGs) with FDR <1%. Schematic of RNA-seq analysis strategy, with “long-growing” (8GW & 12GW) vs. “short-growing” (Neo to 72YO). 20GW #1 and #2 are only provided for a visual comparison and were not included in the differential analysis due to having “variable” growth. **B**) Gene Set Enrichment Analysis (GSEA) for curated genes from developmentally-regulated rodent datasets (orthologs) in long- and short-growing Fib-iNs. Long= genes upregulated in long-growing iNs; short= genes upregulated in short-growing iNs; Up= genes upregulated in rodent developmental datasets, down= genes downregulated in rodent developmental datasets. FDR <0.05 and consistent (red or blue boxes), FDR >0.05 (white boxes), FDR <0.05 but inconsistent (grey boxes). **C**) GSEA for curated gene lists from regenerative rodent datasets (orthologs) in long and short growing Fib-iNs. Up= genes upregulated in regeneration, down= genes downregulated in regeneration. FDR are as in B. **D**) Pearson correlation values and p-values of 6 most significant eigengenes and traits from WGCNA analysis. **E**) Significant Metascape term enrichment in long-growing modules. **F**) Schematic of screen paradigm for overexpression or knockdown of specific genes in Fib-iNs. **G**) Pearson correlation value (r) of genes from the 6 key modules plotted against their respective RNA-seq log2foldchange. Positive r value indicates positively correlated with short-growing phenotype, negative r value indicates positively correlation with long-growing phenotype. Positive log2foldchange indicates gene enrichment in short-growing Fib-iN ages. Negative log2foldchange indicates gene enrichment in long-growing Fib-iN ages. Red dots = genes screened, grey dots = genes not screened. **H**) Z-score of genes tested in 36YO and 8GW Fib-iNs in primary screen. Shaded grey area is 1 standard deviation in each direction. Both red and blue dotted genes were tested in secondary screen, with only red dots successfully reproducing primary screen results. **I**) Secondary screen of 5 primary screen hits, showing the significant genes’ total neurite length normalized to its appropriate control (ctrl; plasmid overexpression (GFP O/E) or scrambled (Scr) shRNA (shRNA)). N ≥ 3 different inductions, n ≥ 50 cells per induction. Student T-test. Mean ± SEM. **J**) Table summary showing secondary screen gene hits, expression trend in human cortex during development (PsychENCODE), knockdown (KD) or overexpression (O/E), age of Fib-iN tested, resulting growth phenotype in the tested Fib-iN age, and WCGNA module. ns = not significant *p<0.05, ***p<0.001.

We next used a Weighted Gene Co-expression Network Analysis (WGCNA) to identify modules of genes that behaved similarly, correlated them to phenotypes (long-growing versus short-growing), and identified significant modules (*50, 51*). Out of the 216 co-expression modules generated, 62 were significantly correlated with a neurite growth phenotype in our Fib-iNs (Fig. 3D, S3C, Table 3-4). We used the top 6 modules with R≥0.89 (2 enriched in short growing, and 4 enriched in long growing) and analyzed the genes within each of these modules further (Fig. 3D, Table 4; https://adsnpheno.shinyapps.io/axonGrowthProject/). In concordance with our phenotypic characterization, long-growing modules enriched for Metascape (*52*) terms associated with axon development and many cytoskeleton reorganizing pathways, suggesting the reorganization of the cellular cytoskeleton structure may be necessary for promoting neurite elongation (Fig. 3E). Short-growing modules enriched for terms associated with receptor signaling and energy generation pathways suggesting that short-growing Fib-iNs may spend more energy responding to the environment (Fig. S3D).

To test the hypothesis that developmentally-regulated genes in human neurons could be candidates for modulating axon growth, we selected a short list of genes to screen for their ability to modify neurite outgrowth in Fib-iNs (Table 5). Using the genes in the selected 6 modules and taking into account the combination of the Pearson correlation score, the fold-change in DEGs from the Fib-iN RNA-seq, and whether the genes were also correctly developmentally-regulated in human brain (*32*), we chose 50 genes to attempt to knockdown (KD) or overexpress (OE) using lentivirus in the appropriate age of Fib-iNs (Fig. 3F-G, S3E). Of the 50 genes tested, 19 genes successfully transduced in our primary screen, and of those 19, 5 genes (26%) significantly modulated neurite growth in the correct direction in either the long-growing or short-growing Fib-iNs, as determined by z-scores (Fig. 3H). In a secondary screen, modulation of 3 of these genes reproducibly significantly modulated neurite growth in the direction predicted from their developmental regulation, including leucine rich repeat kinase 2 (*LRRK2*; coral module), mir9_3p (*MIR9_3HG* was enriched in coral module and mir9_3p regulated many genes in the dark green module), and AT-rich interaction domain 1A (*ARID1A*; darkolivegreen1 module; Fig. 3I-J). *LRRK2* is the most commonly mutated gene in familial Parkinson’s Disease (*53, 54*). Previous studies have found that *LRKK2* OE leads to shortening of neurites, and *LRKK2* knockout leads to increased neurite outgrowth (*54*), similarly confirmed in our results showing that its KD in adult Fib-iNs (36YO) leads to increased neurite outgrowth (Fig. 3I-J). MicroRNA-9 (miRNA-9) is one of the most abundant miRNAs in the brain and has been shown to decrease axon growth in E17 rodent neurons and play critical roles in dendritic development, synaptic transmission, and axon branching (*55–57*). We confirmed a similar role for mir9_3p in human Fib-iNs, where it is upregulated in short-growing neurons, and its overexpression in 8GW neurons leads to reduced neurite growth (Fig. 3I-J). Thus, these data validate the use of developmentally-regulated genes in humans to identify regulators of neurite growth.

ARID1A, also known as Brahma-related associated factor 250a (BAF250a), is a DNA-binding subunit of the canonical mammalian SWI/SNF complex, otherwise known as the canonical BAF complex, which preferentially interacts with enhancers. This ATP-dependent chromatin remodeling complex generally functions as an activator of transcription, with critical roles in regulating chromatin changes during development and in cancer (*58*). Loss-of-function mutations in *ARID1A* can lead to Coffin-Siris Syndrome, a neurodevelopmental disorder with symptoms such as intellectual disability, growth impairment, hypotonia, and structural CNS abnormalities (*59*), but *ARID1A* is most studied for its role in cancer (*60*). Interestingly, both loss- or gain-of-function mutations in *ARID1A* can lead to a wide-range of cancers depending on cellular context, and affects pathways including the WNT, P21, PI3K/AKT, YAP/TAZ, IL-6/STAT3, and IFN pathways (*60*). In the human brain, *ARID1A* mRNA levels decrease during human cortical development (Fig. 4A) (*32*). To determine if ARID1A protein levels were similarly regulated in human development *in vivo*, we assessed the expression of ARID1A and BCL11B, a marker of deep cortical layers, in 16GW, 21GW, and 42YO human cortices. At 16GW, ARID1A protein is present in most cells of the cortical plate. By 21GW, ARID1A expression becomes more spatially restricted within the cortical plate and is enriched in a subset of BCL11B+ neurons. In the adult brain, ARID1A protein levels are dramatically downregulated in the cortex and are lowly expressed across all layers of the cortex (Fig. 4B). These results confirm both published RNA-seq results from human cortical development (Fig. 4A; (*32*)), and our Fib-iN RNA-seq results (Fig. 4C), demonstrating ARID1A is downregulated across development and aging.

**Figure 4:**
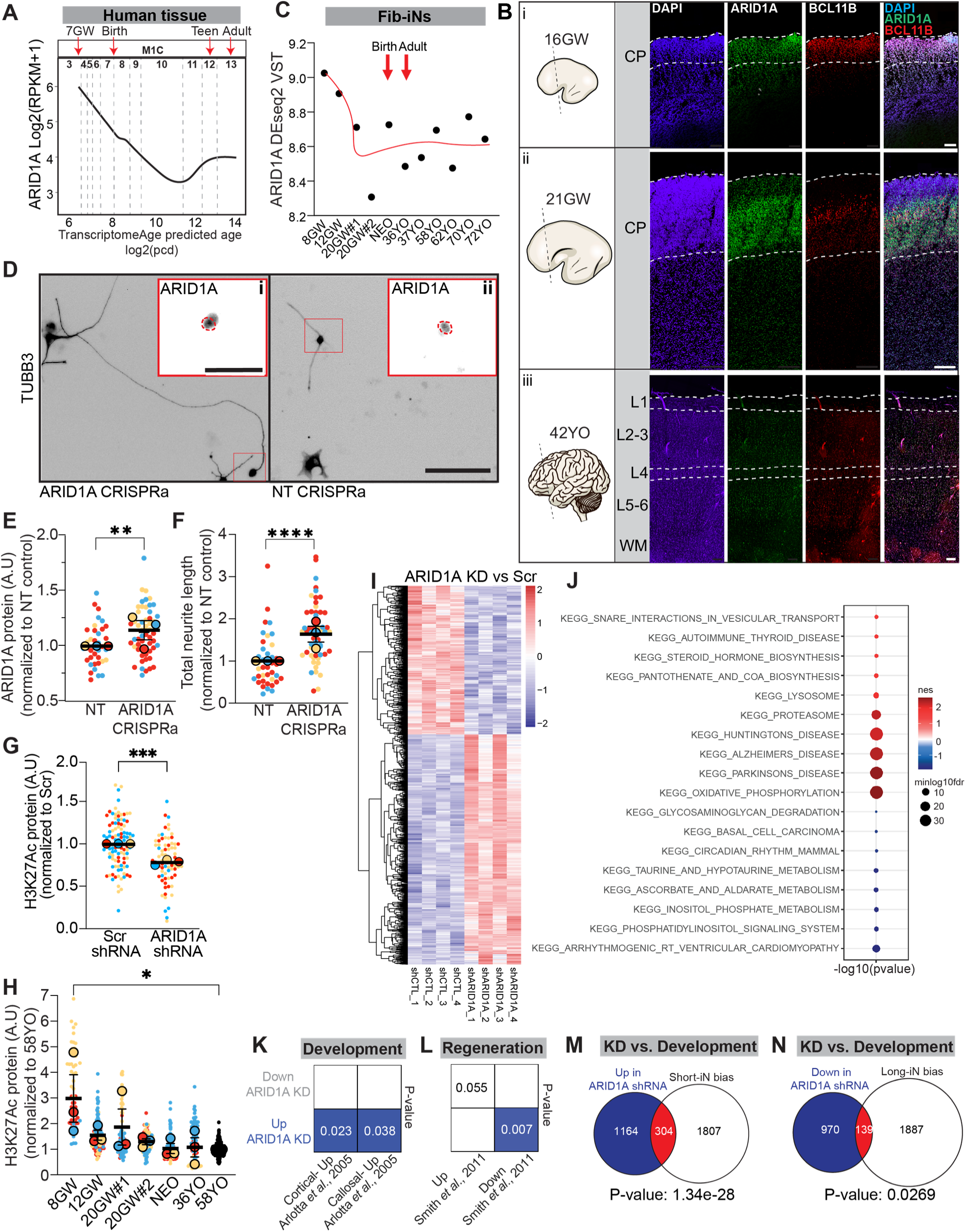
ARID1A is a developmentally-regulated driver of human neurite growth. **A**) ARID1A transcript levels (Log2(RPKM+1) across development in human M1C (PsychENCODE). Post conception day (PCD). Time is displayed in epochs. **B**) Immunostaining of ARID1A in 16GW (i), 21GW (ii) and 42YO (iii) human prefrontal cortex shows colocalization of ARID1A and deep layer neurons (BCL11B). Layer (L) 1-6, CP= cortical plate, WM= white matter. Scale bar, 200 μm. **C**) Average ARID1A transcript expression (DEseq2 VST) across all Fib-iN ages. **D**) Representative images of *ARID1A* CRISPRa (left) and NT (right) control in 36YO Fib-iN, stained for neuronal identity (TUBB3), and ARID1A protein. Scale bar, 100 μm (D), 50 μm (Di, ii). Red circle = nuclei. **E-F**) Quantification of ARID1A protein levels (**E**) with CRISPRa for 7 days in 36YO Fib-iN, and neurite length (**F**; normalized to NT control). Student t-test on averages of replicates; Mean ± SEM. N ≥ 3 different inductions, n=20 cells per induction. **G-H**) Superplot quantification of immunostained H3K27acetylation (H3K27Ac) protein levels **G**) 7-days post ARID1A or Scr shRNA transduction in 8GW Fib-iNs (normalized to Scr shRNA of each experiment), and **H**) across development (normalized to the 58YO of each experiment). N=3 different inductions, n=30 cells per induction. **I**) 1,408 DEGs (FDR <1%) were identified in RNA-seq of ARID1A shRNA vs. Scr shRNA 7 days post-transduction in 8GW Fib-iNs. **J**) Top KEGG pathways significantly enriched in ARID1A shRNA KD as compared to Scr ctrl (red: upregulated with KD, blue: downregulated with KD). **K-L**) Comparison of transcripts upregulated and downregulated in ARID1A KD 8GW Fib-iNs against the same (**K**) developmental and (**L**) regeneration rodent datasets from Fig. 3C-D reveals a similarity of genes upregulated in ARID1A KD and upregulated with development. Up= genes upregulated in rodent datasets; down= genes downregulated in rodent datasets. FDR of <0.05. **M**) Venn diagram revealing the overlap of genes upregulated with ARID1A KD in 8GW Fib-iNs to genes upregulated in short-growing Fib-iNs (Neo-72YO). Fisher’s exact test. **N**) Genes downregulated with ARID1A KD in 8GW Fib-iNs had greater overlap with genes upregulated in long-growing Fib-iNs (8-12GW). Fisher’s exact test. Arbitrary units (A.U.). *p<0.05 **p<0.01 ***p<0.001 ****p<0.0001

Since we found that *ARID1A* KD in long-growing Fib-iNs decreases neurite outgrowth in our screen (Fig. 3I-J, S4A-B), we asked if increasing ARID1A protein levels would be sufficient to increase neurite outgrowth in short-growing Fib-iNs. As ARID1A is a large 250kD protein, we selected CRISPR activation (CRISPRa) to activate its expression in 36YO neurons (short-growing), where a guide RNA (gRNA) localizes an enzymatically dead CRISPR/Cas9 and three transcriptional activators (VP64, HSF1, p65) to the *ARID1A* promoter (OriGene; Fig. S4C). Fib-iNs were transduced with two viruses: one containing a transcriptional enhancer with no reporter, and another containing the gRNA, deadCas9, and GFP (Fig. S4C). We found that GFP+ Fib-iNs transduced with gRNAs targeting *ARID1A* had increased levels of ARID1A protein as compared to the non-targeting gRNA (Fig. 4D-E). Further, we found increased ARID1A protein led to an increase in neurite outgrowth of 36YO Fib-iNs (Fig. 4D, 4F). Thus, taken together, these results demonstrate that modulation of ARID1A protein levels can drive changes in human neurite growth.

As part of the BAF complex, ARID1A primarily targets enhancers thus increasing accessibility for transcription factor binding (*61, 62*). H3K27acetylation (H3K27ac) is a common epigenetic modification associated with the ARID1A-containing BAF complex (*63–66*). Rodent dorsal root ganglion neurons have increased global H3K27ac levels following enriched environment conditions which increased axon regeneration, connecting H3K27ac histone modifications with axon growth potential (*67–69*). KD or loss of ARID1A leads to decreased H3K27ac, resulting in changes in the chromatin architecture (*61, 63, 65, 66, 70*). Thus, we measured the levels of H3K27ac in 8GW Fib-iNs treated with scrambled or *ARID1A*-targeting shRNA. We found reduced levels of H3K27ac with *ARID1A* KD compared to a scrambled control (Fig. 4G). Further, immunostaining of Fib-iNs across developmental ages revealed 8GW as having the highest H3K27ac signal, with a decrease during development and aging (Fig. 4H). These data suggest ARID1A, a developmentally-regulated protein in human neurons, may promote neurite growth at the chromatin level by increasing levels of H3K27ac, thus controlling transcription of downstream targets.

To identify how modifications in ARID1A affect downstream genes and drive changes in neurite outgrowth in Fib-iNs, we performed RNAseq on 8GW Fib-iNs transduced with *ARID1A* shRNA or a scrambled control for 7 days (Fig. S4D-F). We identified 1,408 differentially expressed genes between conditions (FDR<1%; Fig. 4I, Table 6). KEGG terms such as Alzheimer’s, Huntington’s, and Parkinson’s disease were upregulated upon ARID1A KD in 8GW Fib-iNs, whereas KEGG terms for various types of metabolism and phosphatidylinositol signaling were downregulated upon ARID1A KD (Fig. 4J). Excitingly, when we compared transcriptomic changes to previously published rodent studies as in Fig. 3B-C, we found that ARID1A KD shifted the 8GW Fib-iN transcriptome to increase the expression of genes upregulated in cortical and callosal rodent neurons during development, and genes downregulated in rodent PNS regenerating neurons (Fig. 4K-L). Further, when we compared DEGs from ARID1A KD in 8GW Fib-iNs to transcripts upregulated in long-growing and short-growing Fib-iNs, we found that ARID1A KD shifted the 8GW transcriptome to be similar to short-growing Fib-iNs, with significant overlap between genes upregulated upon KD and short-growing biased genes, and between genes downregulated upon KD and long-growing biased genes (Fig. 4M-N). Despite *ARID1A* mRNA levels being only ∼2 fold with the KD (Fig. S4F), this limited change resulted in a reduced neurite growth phenotype mirrored by a shift of the transcriptomic landscape towards that of short-growing Fib-iNs.

Taken together, here we provide an age-maintained and species-specific model for studying neurite outgrowth and identify an *in vitro* developmental switch in human intrinsic neurite growth ability. We find that direct reprogramming can be used to maintain prenatal age in the resulting neurons. We present a novel transcriptional dataset for human neurons with increased or decreased neurite growth abilities. Finally, we identify ARID1A, a DNA-binding subunit of the cBAF nucleosome remodeling complex, as a developmentally-regulated target which can be modified to enhance human neurite outgrowth.

In the embryonic mouse, *Arid1a* deletion in cortical progenitor cells led to corpus callosum agenesis and mistargeting of intracortical axons (*71*), whereas *Arid1a* deletion in the neural progenitors of subplate neurons, the first neurons made in the cortex which pioneer the first axon tracts (*72*), led to decreased growth of descending subplate pioneer axons and loss of subsequent co-fasciculation (*71*). In the adult mouse, *Arid1a* deletion in retinal ganglion cells prior to optic nerve crush had no effect on regeneration (*73*). Indeed, our findings in human Fib-iNs would suggest the opposite strategy in the adult CNS, using *Arid1a* overexpression to increase regeneration. In excitatory human neurons derived from embryonic stem cells, *ARID1A* knockout results in decreased dendritic length and reduced synaptic markers (*74*), though only dendritic growth specifically was measured. Taken together, these and other studies have established a role for ARID1A in pioneer axon growth, dendritic complexity, and synaptic density.

Importantly, identification of ARID1A as a neurite growth regulator implicates chromatin modifiers as important for neurite growth and the lifelong reduction in CNS regenerative abilities, as previously suggested by other studies (*75–79*). However, what drives these chromatin modifications during development is yet unknown. As we find that environmentally naïve human neurons that have never seen another brain region or brain cell demonstrate a loss of intrinsic neurite growth ability similar to what has been seen in rodent CNS neurons, this suggests that an intrinsic clock may drive these changes. Whether these changes are initiated in the body through systemic signals during specific stages of development or the passage of time itself is yet to be determined.

## Acknowledgments

General: We thank Jake Kirkland and Jeffrey Goldberg for comments on the manuscript; the UW-Madison Flow Cytometry core (P30 CA014520 and 1S10RR025483-01); We thank The Birth Defects Research Laboratory for human brain tissue, supported by NIH award number 5R24HD000836 from the Eunice Kennedy Shriver National Institute of Child Health and Human Development; D. Gamm and A. Bhattacharyya for patient fibroblast samples and iPSC lines; J. Mertens for assistance with the direct reprogramming protocol; and members of the Moore lab and UW-Madison community for their input. Artwork was created on Biorender.

## Funding

Hilldale-Holstrom Fellowship (to EANT)

National Institutes of Health grant 1R21NS111192 (to DLM) National Institutes of Health grant F30NS122478-01 (to BPL)

National Institutes of Health grant T32GM140935 (to UW-Madison Medical Scientist Training Program, BPL)

National Institutes of Health grant T32AG052374 (to RJL and BBT) National Institutes of Health grant R35GM142395 (to BAB)

## Author contributions

Conceptualization: BPL, EANT, DLM

Investigation: BPL, EANT, KR, ZPA, PCK, ERP, RR, AMMS, RJL

Analysis: BPL, EANT, KR, BAB

Methodology: EANT, DLM

Software: SS, DW, BAB, BBT, CSM

Funding acquisition: DLM, BAB

Resources: AB

Supervision: BPL, DLM

Visualization: BPL, EANT, DLM

Writing – original draft: BPL, DLM

Writing – review & editing: BPL, EANT, DLM, AMMS, AB, BAB

**Competing interests:** Authors declare that they have no competing interests.

**Data and materials availability:** Processed data are found in the tables. Raw RNA sequencing datafiles found at dbGaP (phs003285.v1.p1) http://www.ncbi.nlm.nih.gov/projects/gap/cgi-bin/study.cgi?study_id=phs003285.v1.p1

## Supplementary Materials

Materials and Methods

Figs. S1-S4

## Tables

Table 1: Original fibroblasts used for experiments.

Table 2: Fib-iN RNA-seq DEG VSTs across all the ages and experimental replicates.

Table 3: WGCNA analysis of all the genes from Fib-iN RNA-seq and their eigengenes.

Table 4: All the genes within the 6 selected eigengenes.

Table 5: Details on genes tested in Fib-iN gene screen.

Table 6: ARID1A shRNA knockdown vs. scrambled control shRNA DEGs.

## Notes

### Competing Interest Statement

The authors have declared no competing interest.

