## Supplement for "Age-maintained human neurons demonstrate a developmental loss of intrinsic neurite growth ability"

**The PDF file includes:**

Materials and Methods
Figs. S1-S4

### Materials and Methods

Cell Culture

**Fibroblast Culture**

Primary human fibroblasts were obtained from the Coriell Institute Cell Repository, A. Bhattacharyya (University of Wisconsin-Madison), and D. Gamm (University of Wisconsin-Madison) (Table 1). Untransduced fibroblasts were cultured in DMEM containing 10% fetal bovine serum (Gibco #10437028). Following lentiviral transduction, fibroblasts were cultured in Tetracycline-Free Medium (TFM), based in DMEM containing 15% tetracycline-free FBS (Fisher Scientific #631106), and 1X nonessential amino acids (Thermo Fisher Scientific #11140050). All fibroblasts were passaged using 0.25% Trypsin-EDTA (Gibco #25200072) for 3 min at 37ºC and 5% $CO_2$.

**Lentiviral Particle Production and HEK 293FT Cell Culture**

For plasmids LV EtO (Addgene #84776), XTP N2AA (Addgene #84777), and pEIGW-hKLF4 (Addgene #100131): HEK 293FT cells were cultured in DMEM containing L-glutamine, 10% fetal bovine serum, and NEAA. Cells were passaged using TrypLE Express. Lentiviral particles were generated by transient co-transfection of HEK 293FT cells with a three-plasmid system using polyethylenimine (PEI; Polysciences, Inc.) and packaging plasmids psPAX2 and pMD2.G. 24 hr after seeding HEK 293FT cells into 10 $cm^2$ BioLite Cell Culture Dishes ($2x10^6$ cells/dish), plasmids and PEI were added to basal DMEM and incubated for 10 min at room temperature (RT). Transfection solution was then added to cells and allowed to incubate at 37ºC and 5% $CO_2$. Lentiviral particles were harvested from the supernatant 2 days after transfection. The supernatant was sterile-filtered (0.22 µm), ultracentrifuged for 2 hr at 175,000 x *g* at 4ºC, and the pellet was resuspended in cold DPBS. Viral particles were titered in human fibroblasts using the antibiotics puromycin and G418 for plasmids XTP N2AA and LV EtO, respectively.

For gene screen plasmids (Table 5) HEK 293FT cells were transfected with third-generation packaging plasmids pREV, pVSVG, pMDL, using Lipofectamine 3000 (Thermo Fisher Scientific #L3000008). Cells were incubated with transfection solution in Opti-MEM for 4-6 hr at 37°C before addition of DMEM + GlutaMAX (10% FBS, 1% NEAA, 0.3% L-Glutamine). Viral particles were harvested as described above.

**Detection of shRNA-mediated knockdown of ARID1A**

Knockdown of ARID1A was detected using HEK 293FT cells, which were transfected with a pLKO.1 lentiviral vector harboring three shRNAs against ARID1A (Table 5) or a scramble control with Lipofectamine 3000 in Opti-MEM. Media was refreshed after 5 hr. Cells were incubated at 37°C for 7 days and collected for detection using western blot analysis.

**Direct Conversion of Adult Human Fibroblasts into iNs**

Following viral transduction, fibroblasts were passaged at least five times with continuous antibiotic selection. To initiate conversion, postnatal fibroblasts were plated at 500K/6-well. After 24 hr, three 6-wells were then pooled together, resulting in ~1.5 million/6-well. Gestational fibroblasts were pooled at 1 million/6-well. For every conversion experiment, the 36YO iN was added as a normalization standard to control for batch effects. 24 hr after pooling, iN conversion medium[14] was added. Medium was refreshed every third day. After 4 weeks of conversion, iNs were gelatin-sorted (see below; [80]) and plated onto poly-L-lysine (PLL, Sigma-Aldrich #P6282) or poly-L-ornithine (PLO, Sigma-Aldrich #P3655) and laminin (Sigma-Aldrich #L2020)-coated ibidi µ-wells in axon supportive medium (ASM) based in Neurobasal containing L-glutamine, N-acetyl-L-cysteine (5 µg/mL; Sigma-Aldrich), B27, N2 (both 1X; Gibco), Forskolin (5 µM; LC Laboratories), BDNF, GDNF (both 20 ng/mL; R&D Systems), natural mouse protein laminin (1 µg/mL; Life Technologies), dibutyryl cAMP (500 µg/mL; Sigma Aldrich), and triiodo-L-thyronine (40 pg/mL; Sigma-Aldrich). After 48 hr post-gelatin sort, Fib-iNs were fixed for ICC.

**Gelatin-based sorting of iNs**

After 4 weeks into induction, iNs were trypsinized using TrypLE and Y-27632 (10 µM; ApexBio) for 45 min at 37ºC and 5% $CO_2$, then quenched in 5% Knockout Serum (KOSR; Thermo Fisher Scientific) in Cyto solution (Myo-inositol, polyvinylalcohol (Sigma-Aldrich)) in PBS and distilled water[14]. Cells were centrifuged at 1,200 x *g* for 5 min and resuspended in TFM. iNs were plated on 0.1% gelatin-coated dishes[80] and incubated for 2 hr at 37ºC. The iN suspension was then centrifuged and plated on PLO or PLL/laminin-coated ibidi µ-slides in ASM.

**Calcein/EI live/dead assay**

To identify the optimal medium for supporting iN viability and neurite outgrowth, iNs were gelatin-sorted after 4 weeks into induction and plated in the following mediums: Neurobasal only (NB), BrainPhys only (BP), Mertens in Neurobasal (M/NB), Mertens in BrainPhys (M/BP), and ASM. Both M/NB and M/BP contained N2, B27 (both 1X; Gibco), BDNF, GDNF (both 20 ng/mL; R&D Systems), natural mouse protein laminin (1 µg/mL; Life Technologies), and dibutyryl cAMP (500 µg/mL; Sigma Aldrich). After 48 hr post-gelatin sort, iN viability in each medium was assessed using a LIVE/DEAD Viability/Cytotoxicity Kit (Invitrogen #L3224) that uses Calcein dye to label live cells and EI dye to label dead cells. Both dyes were diluted 1:1000 in the corresponding basal medium of each medium condition, prewarmed, and added directly to the cells. The cells were incubated with the dyes for 10 min at 37ºC and then immediately imaged.

**Human iPSC culture and reprogramming**

Human induced pluripotent stem cells were generated by the University of Wisconsin-Madison Stem Cell Core. 8GW and 12GW fibroblasts were reprogrammed with the Sendai method (SeV), neonatal fibroblasts were reprogrammed with the episomal method, and 36YO and 72YO fibroblasts were reprogrammed with the retroviral method.

<u>Retroviral iPSC reprogramming:</u>

Fibroblasts were isolated from skin biopsies, cultured in fibroblast media, and reprogrammed by transduction of the following retroviral vectors: pMXs-hOCT3/4 (Addgene #17217), pMXs-hSOX2 (Addgene #17218), pMXs-hKLF4 (Addgene #17219), and pMXs-hc-MYC (Addgene #17220). Vectors were produced by transient transfection of HEK 293T cells with a packaging system that includes Gag-Pol and CMV-VSVG. Cells were plated and cultured with MEF feeder cells, fed with hESCM (human ESC media: DMEM/F12 supplemented with 20% KOSR, 0.1 mM NEAA, 2 mM GlutaMAX, 0.1 mM BME, and 12.5 ng/mL human bFGF) the first week and MEF-conditioned hESCM afterwards. iPSC colonies were manually picked between day 14-28 post-transfection. Cells were expanded and banked on fresh MEF plates with hESCM.

<u>Episomal iPSC reprogramming:</u>

Fibroblasts were isolated from skin biopsies, cultured in fibroblast media, and reprogrammed by electroporation delivery of the episomal vectors: pCXLE-hOCT3/4-

shp53-F (Addgene #27077), pCXLE-hSK (Addgene #27078) and pCXLE-hUL (Addgene #27080). 3 µg of each episomal vector were delivered into $1x10^6$ fibroblast cells using an Amaxa 4d nucleofector. After electroporation, cells were plated and cultured with MEF feeder cells in a low oxygen incubator (5% $O_2$). Cells were fed with hESCM the first week and MEF-conditioned hESCM afterwards. iPSC colonies were manually picked between day 14-28 post-transfection. Following expansion, cells were transferred from MEFs to Matrigel and cultured with mTeSR1 medium (STEMCELL Technologies) for banking.

SeV iPSC reprogramming:

Fibroblasts were isolated from skin biopsies, cultured in fibroblast media, and reprogrammed to iPSCs using the CytoTune-iPS 2.0 Sendai Reprogramming Kit (Thermo Fisher Scientific). $6x10^5$ cells were transduced with Sendai virus at MOI = 5:5:3 (KOS:c-Myc:Klf4). Cells were fed with hESCM for the first week and MEF-conditioned hESCM afterwards. iPSC colonies were manually picked between day 14-28 post-transfection. Emerged clones were picked, expanded, and banked on Matrigel-coated plates using mTeSR medium (STEMCELL Technologies).

Characterization:

Expression of stem cell markers was evaluated by immunostaining. iPSCs were fixed with 4% PFA, permeabilized with 0.1% Triton X-100, and blocked with 10% donkey serum. Primary antibodies against SOX2 (1:1000, R&D # AF2018), NANOG (1:200, Stemgent, 09-0020,), and OCT4 (1:1000, Santa Cruz Biotechnology # SC5279) were incubated overnight at 4°C. After several washes, cells were incubated with the following secondary antibodies: Cy5 donkey anti-rabbit IgG (1:500), 488 donkey anti-mouse IgG (1:500) and Cy3 donkey anti-goat IgG (1:500). Nuclei were stained with Hoechst 33342 (1:1000, Thermo Fisher Scientific # H3570,) Images were captured using a Nikon A1 laser confocal microscope and analyzed using a Perkin Elmer Operetta High Content Analysis System. Used media and live cultures were sent to the WiCell Research Institute for PCR-based mycoplasma testing and G-band karyotyping.

Following reprogramming, iPSCs were grown on Matrigel (Thermo Fisher Scientific #08-774-552), cultured in mTeSR Plus Medium (STEMCELL Technologies #100-0276), and passaged with Versene (Thermo Fisher Scientific #15040066).

**Direct conversion of human iPSCs into induced neurons (iPSC-iNs).**

iPSCs were transduced with LV EtO and XTP N2AA for 4 days, followed by selection with G418 and puromycin for 2 days. Cells were trypsinized with Versene for 3 passages prior to the start of induction in iN conversion medium. To initiate the conversion, iPSCs were plated at 200K/6-well. After 24 hrs, three 6-wells were then pooled together, resulting in 600K/6-well. 24 hrs after pooling, iN conversion medium[14] was added. Medium was refreshed every third day. 5 μM of Cytarabine (Millipore # C3350000) was added to the conversion medium starting at 3 weeks onward. iPSC-iNs were gelatin-sorted at 4 weeks similarly to Fib-iNs as stated in the "gelatin-based sorting" section.

**Fluorescence-activated cell sorting (FACS)**

For RNA-seq (Fib-iN, Fib): After conversion for 4 weeks, Fib-iNs were transduced with a lentiviral hSynapsin-promoter driven mScarlet for 5 days prior to FACS. Cells were sorted based on mScarlet signal using a Baker BioProtect Aria Cell Sorter. mScarlet intensity was binned into 4 bins, with "1" having the highest mScarlet expression, and "neg" having none. 500 cells with the highest mScarlet expression were collected for downstream RNA-seq.

**iN screen with virus**

At 3 weeks into conversion, 36YO or 8GW Fib-iNs were transduced with viral particles for 3 days (overexpression constructs) or 5 days (KD construct) followed by trypsinization and gelatin sorting. Control vector (GFP for overexpression vectors, scramble shRNA for KD plasmids) for every plasmid were included in each respective experiment. Cells were replated in 96-well PLO/Laminin-coated ibidi μ-slides and cultured in ASM. Cells were fixed 48 hr later for analysis.

**CRISPRa transfection**

At 2 weeks into conversion, 36YO Fib-iNs were transfected with 1) pCas-Guide3-GFP-CRISPRa or control pCas9-scrambled-GFP-CRISPRa, and 2) Enhancer vector (OrgiGene Technologies

#GA105467) with Lipofectamine 3000 (Thermo Fisher #L3000008) in Opti-MEM (Invitrogen #11058021) for 3 hr before addition of iN conversion media. 5 days later, cells were trypsinized, gelatin-sorted, replated on PLO/Laminin-coated ibidi μ-slides, and cultured in ASM for 48 hr prior to fixation.

### Immunocytochemistry

#### Staining of Fib-iNs and iPSC-iNs

For all *in vitro* immunostainings, cells were fixed with 4% PFA for 15 min at RT and blocked with 5% donkey serum and 0.2% Triton X-100 based in PBS ($PBS^{++}$). Cells were incubated with primary antibodies diluted in $PBS^{++}$ overnight at 4ºC, washed 3x with PBS, and incubated with secondary antibodies (1:500, Thermo Fisher) for 1 hr and 30 min at RT. Following 3 PBS washes, cells were incubated with Hoechst (1:5000) for 10 min, washed once with PBS, and were then imaged.

Primary Antibodies

| **Antibody** | **Dilution** | **Vendor and catalog #** |
|---|---|---|
| ARID1A | 1:250 | Abcam #ab182560 |
| GABA | 1:500 | Sigma-Aldrich #A2052 |
| GFP | 1:1000 | Aves Labs #GFP-1020 |
| H3K27Ac | 1:1000 | Cell signaling technologies #8173S |
| LMNB1 | 1:1000 | Abcam #ab16048 |
| SOX2 | 1:250 | R&D Systems #AF2018 |
| TUBB3 | 1:1000 | BioLegend #801201 |
| VGLUT1 | 1:1000 | Cedarlane Labs #135302(SY) |
| VIM | 1:1000 | Sigma-Aldrich #AB5733 |
| V5 | 1:500 | Invitrogen #MA5-32053 |

#### Human postmortem samples for immunostaining

Human tissues were collected following the guidelines provided by the University of Wisconsin-Madison Health Sciences Institutional Review Board (IRB) and handled in accordance with ethical guidelines and regulations for the research use of human brain tissue set forth by the NIH (https://oir.nih.gov/sourcebook/ethical-conduct/special-research-considerations/policies-procedures-use-human-fetal-tissue-hft-research-purposes-intramural/policies#acquisition) and the WMA Declaration of Helsinki (https://www.wma.net/policies-post/wma-declaration-of-helsinki-ethical-principles-for-medical-research-involving-human-subjects/). Appropriate informed consent was obtained and all available non-identifying information was recorded for each specimen. All clinical histories, tissue specimens, and histological sections were evaluated to assess for signs of disease, injury, and gross anatomical and histological alterations. No obvious signs of neuropathological alterations were observed in any of the human specimens considered and analyzed in this study.

| **Species** | **Source** | **Age** | **Sex** |
|---|---|---|---|
| Human | Yale School of Medicine | 42 years-old | Male |
| Human | Birth Defects Research Laboratory, University of Washington | 19 postconceptional weeks/ 21 gestational weeks | Male |
| Human | Birth Defects Research Laboratory, University of Washington | 14 postconceptional weeks/ 16 gestational weeks | Female |

**Human tissue processing**

Tissue samples from the brain specimens analyzed in the study were fixed in 4% PFA and incubated in progressive solutions of 10%, 20%, and 30% sucrose. Adult human sections were cut on a Leica VT1000S Vibratome and 50 µm sections were mounted on TOMO® adhesion slides (Matsunami Glass USA #TOM-11/90). Human fetal sections were embedded in OCT, frozen in isopentane-dry ice slurry, and 20 µm sections were cut on a Leica CM1950 Cryostat and mounted on TOMO® adhesion slides (Matsunami Glass USA #TOM-11/90).

**Immunohistochemistry of human sections**

Adult human brain sections were washed in PBS (3 x 5 min) and antigen unmasking performed with the Retriever 2100 (Electron Microscopy Services) using Buffer A (EMS #62706-10). Sections were washed in PBS (2 x 5 min) and incubated in blocking solution (5% normal donkey serum (Jackson ImmunoResearch Laboratories), 0.3% Triton X-100) in PBS for 30 min at RT.

OCT-embedded human fetal brain sections were washed in PBS (3 x 15 min) and incubated in blocking solution for 30 min at RT. Primary antibodies – ARID1A (1:250, Abcam #ab182560), BCL11B (1:500, Abcam #ab18465) – were diluted in blocking solution and incubated with tissue sections for 24 hr at 4°C. Sections were washed with PBS-T (PBS + 0.3% Triton X-100) prior to being incubated with secondary antibodies (1:250, Jackson ImmunoResearch Labs) in blocking solution for 1 hr at RT. Sections were then washed with PBS-T (3 x 5 min), treated with Autofluorescence Eliminator Reagent (Millipore #2160) according to manufacturer instructions, and cover-slipped with Vectashield Plus Antifade Mounting Medium (Vector Laboratories #H-1000).

**Imaging of human sections**

Z-stack, tiled images of human immunohistochemical sections were acquired using a Nikon A1 confocal microscope.

**Western blot analysis**

To collect a soluble protein fraction, cells were lysed in soluble lysis buffer (10 mM Tris HCl, 150 mM NaCl, 0.5 mM EDTA, 0.5% NP-40, 1X protease inhibitor (Sigma #04693124001)) and were resuspended every 10 min for 30 min on ice. Lysed samples were then centrifuged at 20,000 x *g* for 10 min at 4°C, and the supernatant was collected. Total protein extraction was performed by resuspending cell pellets in total lysis buffer (10 mM Tris pH 7.4, 1% Triton X-100, 150 mM NaCl, 10% glycerol, 4% SDS, 1X protease inhibitor, DNAse (1.6 mg/ml)). Lysed samples were sonicated with a probe sonicator for 15 sec on low-power setting. All samples were then quantified using the DC assay (Bio-Rad #5000112) and 20 μg of protein per sample were run through an SDS-PAGE gel. Gels were then transferred onto a PVDF membrane at 100V for 1.5 hr at 4°C. Membranes were then blocked for 30 min at RT in 5% milk made in 1X TBS-T (0.1% Tween-20, 20mM Tris HCl, 150mM NaCl). Primary antibodies (1:1000; ARID1A (Abcam #ab182560), ACTB (BioRad #VMA00048)) were incubated with membranes overnight in 5% milk at 4°C. Membranes were washed the next day (3 x 10 min) in TBS-T. The membranes were then incubated with secondary antibodies (1:3000, BioRad) in 5% milk + TBS-T for 1.5 hr at RT and then washed (3 x 10 min) with TBS-T. Proteins were visualized with a UVP imaging system after treating blots with SuperSignal Femto or Pico (Invitrogen #34095, #34577). Protein expression was quantified using FIJI.

**Imaging/Analysis**

**In vitro imaging**

For all neurite tracing analysis, cells were imaged on a Zeiss epifluorescent microscope with a 10X objective, unless indicated otherwise.

**Analysis of neuronal conversion efficiency and post-sort efficiency (PSE)**

For quantification of neuronal conversion efficiency of iNs, cells were binned into one of four categories based on expression of beta-III tubulin (TUBB3) and vimentin (VIM; for Fib-iN) or SOX2 (for iPSC-iN): iNs ($TUBB3^+/VIM^-$; $TUBB3^+/SOX2^-$), fibroblasts ($TUBB3^-/VIM^+$) or iPSCs ($TUBB3^-/SOX2^+$), fibroblast or iPSC intermediates ($TUBB3^-/VIM^-$; $TUBB3^-/SOX2^-$), and iN intermediates ($TUBB3^+/VIM^+$; $TUBB3^+/SOX2^+$). The total number of cells in each bin was quantified at indicated time points throughout induction and 48 hr following gelatin-based sorting at 4 weeks post-induction (PSE). For each analysis, ten 20X images were analyzed for each technical triplicate, which was repeated across three separate experimental inductions.

**Characterizarion of neuronal subpopulations**

GABAergic iNs (Fib-iN and iPSC-iN) were defined by $GABA^+$ cell bodies and neurites, while glutamatergic iNs were distinguished by $vGLUT1^+$ cell bodies and neurites. Analysis was performed 48 hr after gelatin-based sorting at 4 weeks post-induction. The percentages of $GABA^+$ and $vGLUT1^+$ iNs were calculated based on total number of $TUBB3^+$ neurons. For each analysis, ten 20X images were analyzed for each technical triplicate, which was repeated across three separate inductions.

**Quantification of neurite length and branching**

For quantifying Fib-iN and iPSC-iN neurite outgrowth: iNs fixed 48 hr following gelatin-based sorting at 4 weeks post-induction. Only $GABA^-/TUBB3^+$ iNs with healthy neurites that successfully initiated a neurite length of ≥ 5 μm[3], were analyzed on FIJI. The Longest Neurite and its Branches (LNB; includes only the longest neurite extending from the cell body and its branches) and Total Neurite Length (TNL; includes all neurites extending from the cell body and their branches) were analyzed for each neuron. All neurite lengths were quantified using FIJI. For each age, 50-100 iNs were analyzed for each technical triplicate, which was repeated across three separate inductions. To control for induction-based variability, the 36YO iNs were used as a normalization control in every experiment, unless indicated otherwise. The LNB and TNL of each neuron analyzed were normalized to the respective 36YO sample from each experimental induction.

For Fib-iN gene screen with overexpression or knockdown: iNs were transduced per "iN screen with virus," fixed, and stained as indicated above. iNs ($TUBB3^+/VIM^-$) successfully transduced with virus ($GFP^+$ or $V5^+$) were identified and TNL was quantified. TNL of each virus and age was normalized to the virus and age control of each induction. For each induction, a minimum of 30 cells were quantified for the primary screen (Fig. 3H) and only induction per virus. For the secondary screen, a minimum of 50 cells were quantified (Fig. 3I-J), and 3 separate inductions were performed.

For ARID1A CRISPRa screen: iNs were transfected per "CRISPRa transfection" section, then fixed and stained as indicated above. iNs ($TUBB3^+$) successfully transfected with CRISPRa plasmid ($GFP^+$) were identified and their TNL and ARID1A protein signal via immunostaining were quantified. The TNL of ARID1A CRISPRa cells were normalized to the average TNL of the non-targeting control of each induction. 20 $TUBB3^+/GFP^+$ cells were quantified per condition, 3 separate inductions were performed.

**Analysis of neurite morphology**

Neurons with neurite blebbing were defined by having at least 3 $TUBB3^+$ spherical blebs across that neuron's neurites. The percentage of neurons with neurite blebbing was quantified based on total number of $TUBB3^+$ neurons at 48 hr following the gelatin-based sort at 4 weeks post-induction. Ten 20X images were analyzed for each technical triplicate, which was repeated across three separate inductions.

**LMNB1 immunofluorescence signal quantification**

For quantification of laminB1 (LMNB1) fluorescence intensity: Cells were imaged at 60X on Nikon C2 confocal microscope with the same laser power. All cells for the same induction were imaged within the same day. Samples were blinded prior to analysis on FIJI. Z-stacks were max projected and analyzed with a macro made by our lab in FIJI. Mean A.U. of the measured area was divided by the area to give an A.U./area value for each cell which was normalized to the average A.U./area for the 36YO of every induction. A minimum of 50 cells were quantified per induction, and 3 separate inductions were performed.

For quantification of H3K27Ac: Cells were imaged on Zeiss epifluorescent microscope with a 10X objective with the same light settings. All cells from the same induction were imaged within the same day. Samples were blinded prior to analysis on FIJI. iNs ($TUBB3^+/VIM^-$) were identified and a mask was drawn in the Hoechst channel surrounding the nuclei, then the same mask was applied to the H3K27Ac antibody channel where the raw intensity in A.U and area were collected per cell. Then the A.U/area was calculated per cell and normalized to the 58YO of the same induction. A minimum of 30 cells were quantified per condition per induction, and 3 separate inductions were performed. Same method was applied to ARID1A protein level quantification after immunostaining, with the exception that criteria for identification of the target cell was for the cell to be $TUBB3^+$(iN) and $GFP^+$(CRISPRa). The A.U/area of ARID1A CRISPRa iNs were normalized to the nontargeting control within each induction. A minimum of 20 cells were quantified per induction per condition, and a total of 3 inductions were performed and quantified.

**RNAseq library construction**

For low input bulk RNA-seq, 500 cells were FACS sorted directly in qPCR plates. The 500 cells were subjected to cDNA synthesis using the SMART-Seq® v4 Ultra® Low Input RNA Kit (Clontech cat #634889) according to the manufacturer's protocol. cDNA were quality controlled using high sensitivity D5000 screen tapes on the Agilent Tapestation 4200 before library generation. 150 pg of the amplified cDNA were used to construct cDNA libraries using the Nextera XT DNA Library Preparation Kits (Illumina cat #FC-131-1024 and FC-131-1096) following the manufacturer's protocol. Libraries were quality controlled using high sensitivity D1000 screen tapes on the Agilent Tapestation 4200. Paired-end 150-bp reads were generated on the Illumina Novaseq 6000 platform at the Novogene Corporation.

**RNAseq processing and alignment**

Paired-end 150bp RNA-seq reads were first hardtrimmed to retain bases 9 to 100 (to remove biased sequences due to not-so-random priming on the 5' end, and lower quality sequences on the 3' end) using fastx_trimmer from the fastx toolkit 0.0.13 [http://hannonlab.cshl.edu/fastx_toolkit/index.html]. Hard trimmed reads were then processed for adapter removal and quality trimming using TrimGalore 0.6.6 with parameters -q 15 --length 75 [https://www.bioinformatics.babraham.ac.uk/projects/trim_galore/]. Cleaned up RNA-seq reads were aligned to the hg38 human genome using STAR 2.7.7a[81], with parameters --outFilterMultimapNmax 50 --outFilterIntronMotifs RemoveNoncanonicalUnannotated. Aligned reads were summarized over genes to obtain a gene count matrix using featureCounts from subread 1.5.3[82].

**Genetic PC calculation**

Aligned reads from all independently reprogrammed iN samples derived from the same initial donor were merged using samtools merge (Samtools 0.1.19). Genetic variant identification from RNAseq was performed as recommended by the GATK project [https://gatk.broadinstitute.org/hc/en-us/articles/360035531192-RNAseq-short-variant-discovery-SNPs-Indels-]. Briefly, we used gatk 4.1.9.0[83] to determine genotype at detected sites. Sequentially, each sample was processed with MarkDuplicates, SplitNCigarReads, AddOrReplaceReadGroups, BaseRecalibrator, ApplyBQSR, HaplotypeCaller, VariantAnnotator and VariantFiltration (with options --filter-expression "MQ0 >= 4 && ((MQ0 / (1.0 * DP)) > 0.1)" --filter-name "HARD_TO_VALIDATE" --filter-expression "QUAL < 50 && DP > 5" --filter-name "GATKStandard"), yielding only high confidence variants in a single vcf file per original donor. All sample calls were merged into one file with bcftools merge (--missing-to-ref --apply-filters PASS). Finally, we used plink 1.90beta to prune the data (options --indep-pairwise 50 10 0.1) and calculate principal components (PCs) in the data[84].

**Differential expression analysis**

The read count matrix was loaded into R 3.6.3 for downstream processing. For the multi-donor iN analysis, data was corrected for potential surrogate variables using package SVA 3.34.0[85], for genotype PCs 1-5, as recommended for aging analysis in human data[86] and for batch. No correction was applied for the shRNA analysis since all cells were derived from the same donor. Only transcripts that were expressed in at least 20 of the 29 RNA-seq samples (iN aging dataset) or 4 out of 8 (shARID1A dataset) were retained for downstream analysis. DEseq2 1.26.0[87] was used to perform differential gene expression analysis for (i) genes significantly regulated in long-growing vs. short-growing iNs, and (ii) for age-related expression changes (in 36YO and older). Significant genes were plotted as a heatmap.

**Neuron identification analysis**

We obtained transcriptional profiles of neuron types (Human_Multiple_Cortical_Areas_SMART-seq) from Allen Brain atlas portalmap [https://portal.brain-map.org/explore/classes/nomenclature; access 2021-03-29)[21,22]. We specifically obtained TMM normalized count values from neuronal cells. For each pair of neuronal type from portal map and Fib-iN samples (using normalized data; see above), spearman rank correlation (Rho) was computed. Higher Rho values suggest higher similarity of expression profiles.

**Transposable Element Differential Expression Analysis**

Genes and TEs were counted from STAR aligned BAM files using TETranscripts v2.2.1[88] with the genic gtf and TE gtf options set to "--GTF hg38.refGene.gtf --TE hg38_rmsk_TE_20200804.gtf", respectively. Gene and TE counts from TETranscripts were then loaded into R v4.0.1 for processing. Only transcripts that were expressed in at least 20 of the 29 RNA-seq samples were retained. Technical variation in transcript expression was removed using SVA v3.38.0 and genotype PCs 1-5 from plink (see above). DESeq2 v1.30.1 was used to run differential expression analysis considering both genes and TEs[87]. TEs that were differentially expressed (FDR <5%) were subsetted, and TEs with significant expression changes were plotted as a heatmap.

**Gene Set Enrichment Analysis (GSEA)**

GSEA was used to determine whether (i) axon length related, (ii) aging related or (iii) *ARID1A* knock-down responsive genes showed functional enrichment for categories of interest[89]. Briefly, the DESeq2 moderated t-statistic (column 'stat') was used to rank genes by amplitude of response to the conditions evaluated. We used phenoTest 1.34.0 and qusage 2.20.0 in R 3.6.3 to perform

GSEA, with B = 10000, minGenes = 5, and maxGenes = 7500. We performed 2 sets of GSEA analyses, one on standard gene sets (GO, KEGG) derived from MSigDB v7.3 and one on curated gene sets from the literature[33,42,43,45-49]. Normalized Enrichment Score (NES) was used to determine directionality of enrichment, and the Benjamni-Hochberg corrected p-values were used to determine significance of enrichment.

**Machine learning model (S2I)**

Summarized gene count table and metadata for bulk mRNA-seq data from PsychENCODE[32] were downloaded on 2021-04-15. Data was subsetted to retain M1C and V1C samples and only developmental ages before 3rd trimester, yielding 26 samples ranging 56 to 154 dpc. Count data was corrected for potential surrogate variables using package SVA 3.34.0 in R 3.6.3[85]. To aid in feature selection, DEseq2 1.26.0[87] was used to perform differential gene expression as a function of developmental age, and transcripts with (i) significant (FDR < 5%) changes in both brain regions, (ii) in the same direction during development and (iii) also detected in the Fib-iN RNAseq were retained for downstream analyses (1,669 genes). SVA-corrected counts for the Fib-iN from developmental time points were merged with the filtered/corrected PsychENCODE data[32] to perform joint TMM normalization and computing sample-wise cpm values for all samples using edgeR 3.28.1[90]. Cpm values were then $\log_2$ transformed for downstream analysis. In addition, since PsychENCODE included both male and female samples, sex was also used as a covariate (all used Fib-iN were male). Package caret 6.0-86 was used to train a generalized linear model (GLM) on the PsychENCODE data[32] using leave-one-out cross validation (LOOCV; due to lower sample size) and RMSE optimization. The trained model was then used to predict the gestational age of Fib-iN samples using the same feature set.

**WGCNA analysis**

***Weighted gene co-expression network analysis***

Given the relatively large number of differentially expressed genes that we had identified in our RNA gene expression data, we wanted to utilize pathway analysis to better hone in on potentially key tightly-coordinated genes for our analysis. Our RNA gene expression data had 16,813 genes for 29 samples, 11 different ages. This was used as input gene expression data for the Weighted Gene Co-expression Network Analysis (WGCNA)[50]. We used the suggested soft thresholding power of 30 to build our adjacency matrix. In the end, we constructed gene co-expression networks, which we then clustered into gene co-expression modules. Each of these gene modules contained at least 30 genes. To improve the biological significance of our modules, we applied an additional K-means step[91] based on the number of modules WGCNA had detected, the modular eigengenes (MEs) from our WGCNA modules as the starting centroids, and the starting assignments of genes in the respective modules. Then, we ran this K-means step to convergence, reassigning genes to optimal modules across the iterations. Ultimately, we found over 215 gene modules for our 16,813 genes.

***Enrichment analyses of gene co-expression modules***

Co-expressed genes in the same module are highly likely involved in similar functions and pathways as they share expression dynamic patterns. Thus, gene enrichment analysis has been widely used to identify such functions and pathways in a group of related genes like a gene module. Enrichment p-values were adjusted using the Benjamini-Hochberg (B-H) correction. Given a group of genes (e.g., from a gene co-expression module), we used multiple tools for enrichment analyses, including g:Profiler and Metascape. We have gathered enrichment information from hundreds of different sources for each of our gene modules across the brain regions. Additional R packages were used, such as: ABAEnrichment, WGCNA, Psygenet2r, ClusterProfiler, DOSE, msigdbr. Databases (such as MaayanLab and BaderLab) were consulted to provide more enrichments.

***Association of genes and modules with neurite growth-related phenotypes***

We further associated genes and modules with our key neurite growth-related phenotypes: long-growing, variable, and short-growing neurites. Long-growing Fib-in ages were the 2 averaged samples corresponding to 8 and 12GW, respectively; variable neurites were those 2 samples in 20GW #1 and #2; short-growing Fib-iN ages were Neo, 36YO, 37YO, 58YO, 62YO, 70YO, and 72YO. In addition, we derived further phenotypes like "not long" (in either short or variable growing categories) and "not short" (in either long or variable growing categories). We found the Pearson correlations of each of our modular eigengenes (MEs) and each of our respective genes with each of the neurite growth-related phenotypes. A modular eigengene is a vector with its elements representing the expression levels of input samples and represents the most likely gene expression patterns of modular genes. In essence, an ME typically provides a snapshot of its respective gene co-expression module and is the first principal component of gene expression levels across the samples in that module. We used the moduleTraitCor() and moduleTraitPvalue() functions in WGCNA to then find significantly positively correlated neurite growth-related phenotypes to the modules ($p < 0.05$ and $r > 0$). Furthermore, we identified potential hub genes for each of the modules.

**Statistical analysis**

All non-genomic statistical analyses were performed in GraphPad Prism 9 using tests as indicated in the figure legends. All data were tested for normal distribution using the Shapiro-Wilk normality test. Datasets with a normal distribution underwent parametric tests for significance, whereas datasets without a normal distribution underwent nonparametric tests for significance. An Unpaired Student's t-test was utilized for comparing two groups with normal distribution. The Mann-Whitney test was utilized for comparing two groups with non-normally distributed data. When comparing more than two groups with normally distributed data, a two-way ANOVA was used followed by a post-hoc Tukey's test for group comparisons. The Z-scores were calculated by averaging the normalized fold change for TNL in virus transduced conditions over the control to derive a mean. The standard deviation was then calculated within each age cohort (8GW or 36YO). Z-score value for each gene screened was then calculated by (normalized TNL – mean)/standard deviation. For cell culture experiments, ''N'' represents the number of times the experiment was repeated on different days, ''n'' is equal to the number of cells analyzed.

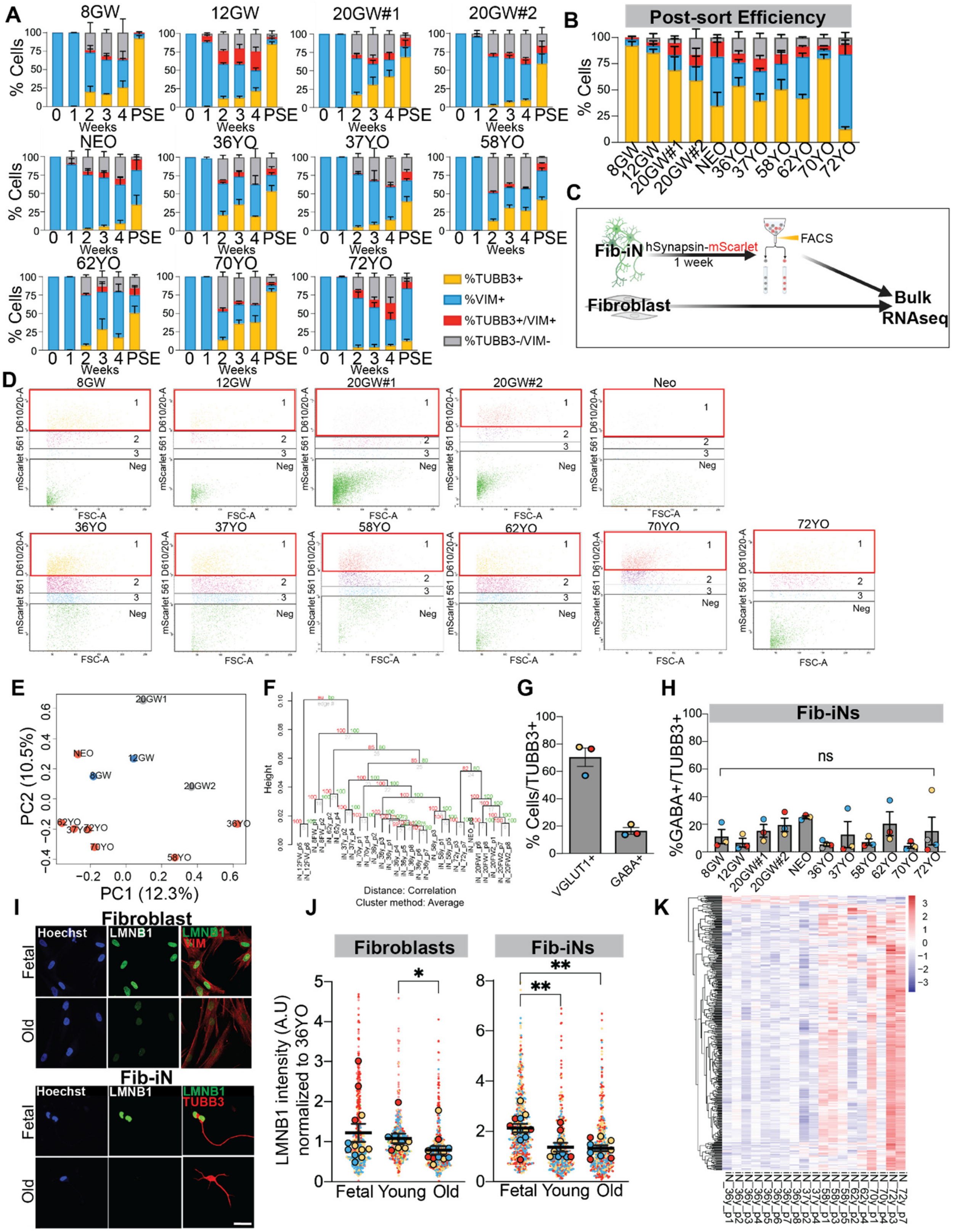


**Fig. S1: Characterization of Fib-iNs.**

A) Quantification of neuronal conversion efficiency of each fibroblast line at 0 weeks (W), 1W, 2W, 3W, 4W, and 48 hours post-sorting (post-sort efficiency; PSE) based on expression of TUBB3 and/or VIM (Fibroblasts (TUBB3-/VIM+; blue), intermediates (TUBB3+/VIM+; red), iNs (TUBB3+/VIM-; yellow), or unnamed (TUBB3-/VIM-). Three separate inductions are represented per age. B) Quantification and comparison of neuronal conversion efficiency 48 hours post-gelatin sort across all ages of Fib-iNs based on expression of TUBB3 and/or VIM. C) Fib-iNs of each age were transduced with a lentivirus encoding a human Synapsin promoter driving mScarlet, sorted using fluorescence-activated cell sorting (FACS), and sent for RNA-seq. Original fibroblast lines were also sorted using FACS and sent for bulk RNA-seq. D) Representative FACS plots of mScarlet+ Fib-iNs from each age. Bin 1 represented the highest intensity of mScarlet expression, and 500 of these cells were collected from each age for bulk RNA-seq. E) Principal component analysis (PCA) of genotypes of samples, derived from RNA-seq of all ages of Fib-iNs. F) Hierarchical clustering of transcriptional profiles of Fib-iNs using pvclust, with pvalues (%), showing consistency between Fib-iN replicates. G) Quantification of glutamatergic (VGLUT1+) and GABAergic (GABA+) 36YO Fib-iNs, demonstrating that iNs produced with this protocol are primarily glutamatergic. Student's t-test, N =3; Mean ± SEM. H) Quantification of GABAergic Fib-iNs across all ages, demonstrating that similarly low percentages of GABA+ iNs generated with this transdifferentiation protocol. One-way ANOVA with post-hoc Tukey's test on averages of replicates; Mean ± SEM. I) Representative images of immunostaining of the nuclear envelope marker, Lamin B1 (LMNB1; green) in fetal (8GW) and old (72YO for fibroblasts, 70YO for Fib-iN) fibroblasts and Fib-iNs. Fibroblasts (VIM+; red), neurons (TUBB3+; red), nuclei (Hoechst). Scale bar, 50 µm. J) Fluorescence intensity of binned ages (Fetal (8GW-20GW), Young (Neo-37YO), Old (58YO-72YO)) were normalized to the 36YO sample within each experiment. Superplots display single cells from each experimental replicate (small dots, colors indicate separate experimental replicates), and the average of each replicate (large dots, color-matched to small dots). One-way ANOVA with post-hoc Tukey's test on averages of replicates; Mean ± SEM. Arbitrary units (A.U.). K) Differential expression analysis of transposable elements (TEs) in adult Fib-iNs (36YO-72YO) reveals 422 TEs upregulated (FDR<5%) as a function of age. ns = not significant, *p<0.05 **p<0.01.

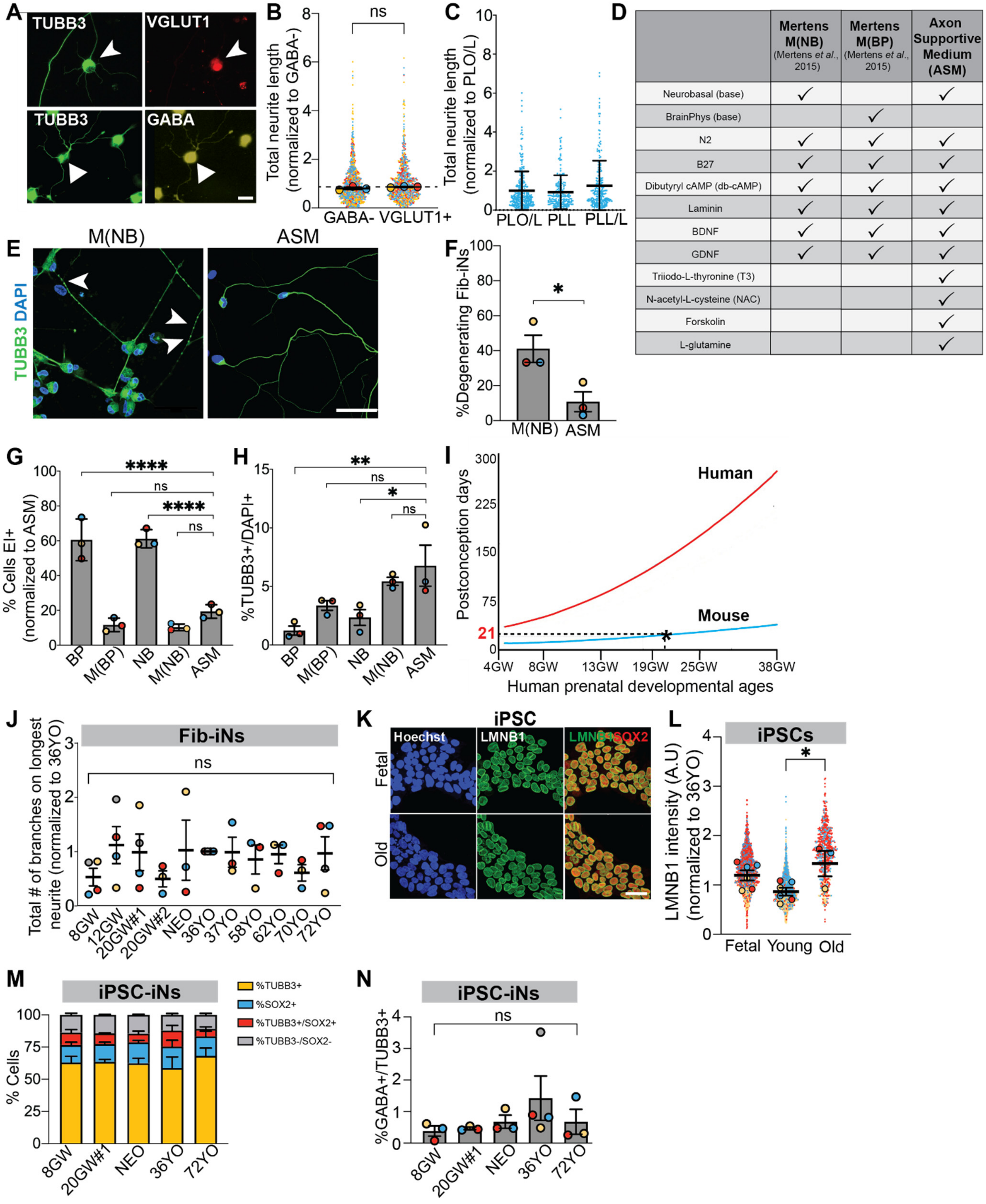


**Fig. S2: Characterization of Fib-iN and iPSC-iN identity, health and neurite growth.**

Examples of 36YO GABAergic (GABA; yellow) and glutamatergic (VGLUT1; red) Fib-iNs (TUBB3; green). Arrowheads point to a VGLUT+ Fib-iN. Triangles point to a GABA+ Fib-iN, based on signal also present in neurites. Scale bar, 50μm. B) Total neurite length quantification of excitatory 36YO Fib-iNs using either GABA- staining or VGLUT1+ staining (normalized to GABA- population; dashed line) reveals no significant difference between staining strategies to detect excitatory neurons. N ≥ 3 different inductions, n ≥ 50 cells. C) Total neurite length quantification of 36YO Fib-iNs cultured on poly-L-ornithine (PLO)/laminin, poly-L-lysine (PLL) only, and PLL/laminin (normalized to the PLO/laminin condition) demonstrates that substrate type does not affect Fib-iN neurite outgrowth. N=1 induction, n=50 cells. D) Ingredient list of previously published neuronal maturation media (*14*) in Neurobasal or BrainPhys (M(BP)) compared to axon supportive medium (ASM) used to culture low density 36YO Fib-iNs post-sort. E) Immunostaining of TUBB3 reveals increased neurite blebbing and degeneration of Fib-iNs cultured at low density in M(NB) in contrast to ASM. Arrowheads indicate blebs. F) Quantification of neurite degeneration, defined as any Fib-iN with three or more distinct blebs across its neurites, of 36YO Fib-iNs cultured at low density in ASM and M(NB). ASM supports healthy, intact neurites at low density. Student t-test on averages of replicates. N ≥ 3 different inductions, n ≥ 50 cells. G) Calcein/ethidium homodimer-1 (EI) live/dead survival assay reveals similar viability of 36YO Fib-iNs 48 hrs post-sorting. (N ≥ 3 inductions, n ≥ 50 cells; ANOVA with post-hoc Tukey's test compared to ASM). H) ASM and M(NB) similarly promote conversion efficiency of 36YO iNs post-sorting in comparison to BP, M(BP), and NB. (N ≥ 3 different inductions, n ≥ 50 cells; ANOVA with post-hoc Tukey's test compared to ASM replicates). I) Human gestational ages compared to mouse development using neurodevelopmental stages (based on *(38)*) reveal human ~21GW as the equivalent of rodent birth (E22/P0), which corresponds to a developmental decrease in central nervous system (CNS) rodent axon growth ability (*4*). J) Branching analysis reveals no differences in total number of branches on the longest neurite across all ages/samples (normalized to 36YO). ANOVA with post-hoc Tukey's test on averages of replicates. N ≥ 3 different inductions, n ≥ 50 cells. K-L) Representative immunostaining images of LMNB1 from fetal (8GW) and old (72YO) iPSCs (SOX2), nuclei (Hoechst). Scale bar, 50μm. L) Quantification of fetal (8GW-12GW), young (Neo-36YO), old (72YO) iPSC LMNB1 protein intensity in arbitrary units (A.U) normalized to the 36YO of each experimental replicate. N ≥ 3 different inductions, n ≥ 50 cells per induction. M) Post-sort efficiency across 8GW, 20GW #1, NEO, 36YO, and 72YO iPSC-iNs show similar rates of iN induction across all the ages (iPSC-iN (TUBB3+/SOX2-), iPSCs (TUBB3-/SOX2+), intermediates (TUBB3+/SOX2+), and undefined (TUBB3-/SOX2-). Three separate inductions are represented per age. N) Quantification of %GABAergic iPSC-iNs following transdifferentiation, revealing no difference between ages. One-way ANOVA with post-hoc Tukey's test on averages of replicates. N $\geq$ 3 different inductions, n $\geq$ 50 cells per induction. Mean ± SEM. ns=not significant *$p<0.05$ **$p<0.01$ ****$p<0.0001$.

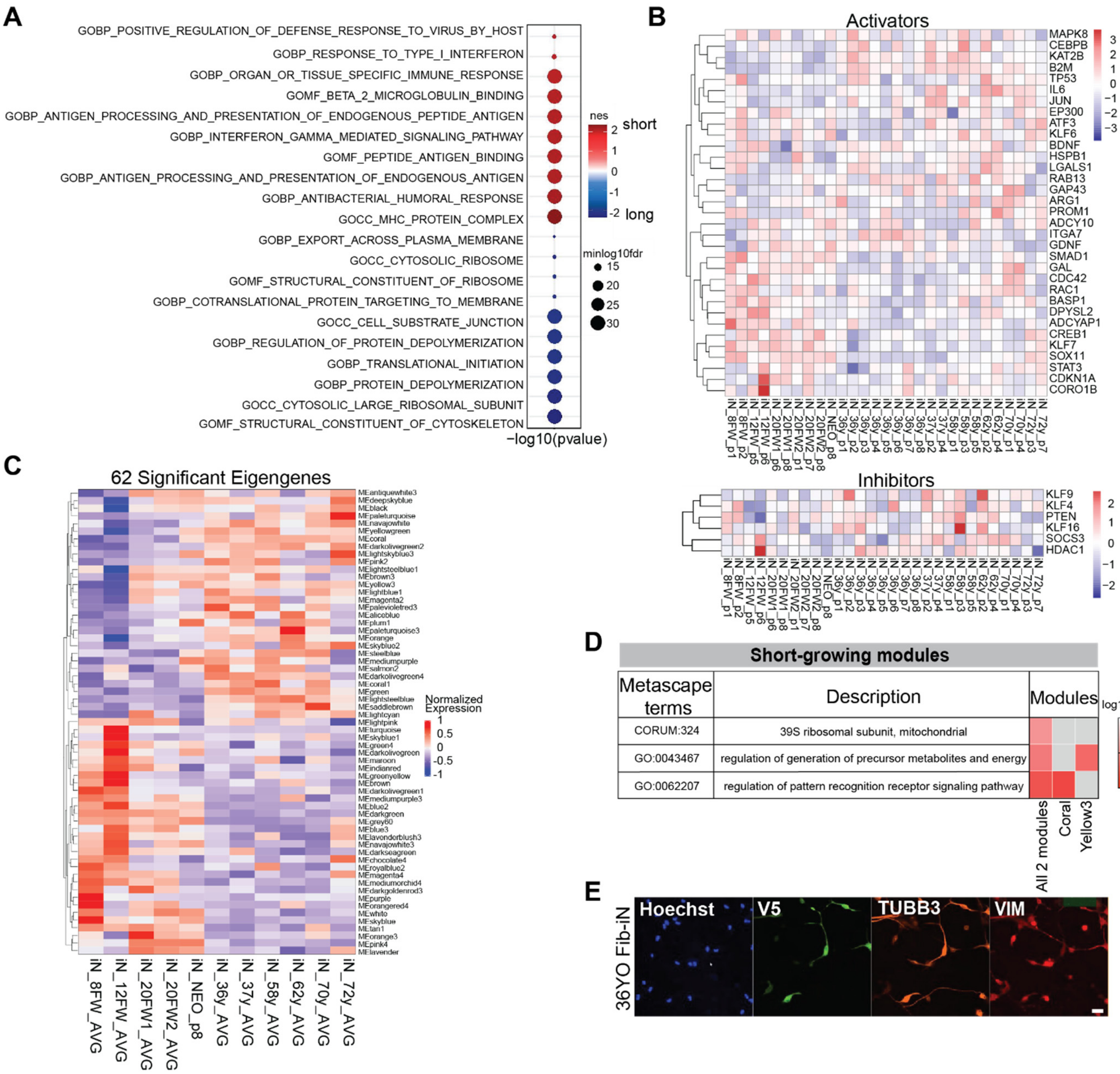


**Fig. S3: Downstream analysis of Fib-iN RNA-seq**

A) Top GSEA terms based on comparison of long-growing and short-growing Fib-iNs (red – more in short; blue – more in long). B) Heatmap of a curated list of the expression (DEseq VST) of previously identified rodent axon growth activators and inhibitors in Fib-iNs over development. C) Heatmap showing 62 significant eigengenes from WGCNA analysis based on the average of replicates from every Fib-iN age, and their relative expression in modules. D) Top significant Metascape term enrichment (Log10(P)) in short-growing modules (coral, yellow3). E) Representative images of 36YO Fib-iNs transduced for 5-days with a V5 control overexpression plasmid. Fib-iNs were immunostained for the V5 epitope tag (transduced, green), TUBB3 (neurons, orange), VIM (fibroblasts; red), and Hoechst (nuclei, blue). Scale bar, 10 μm.

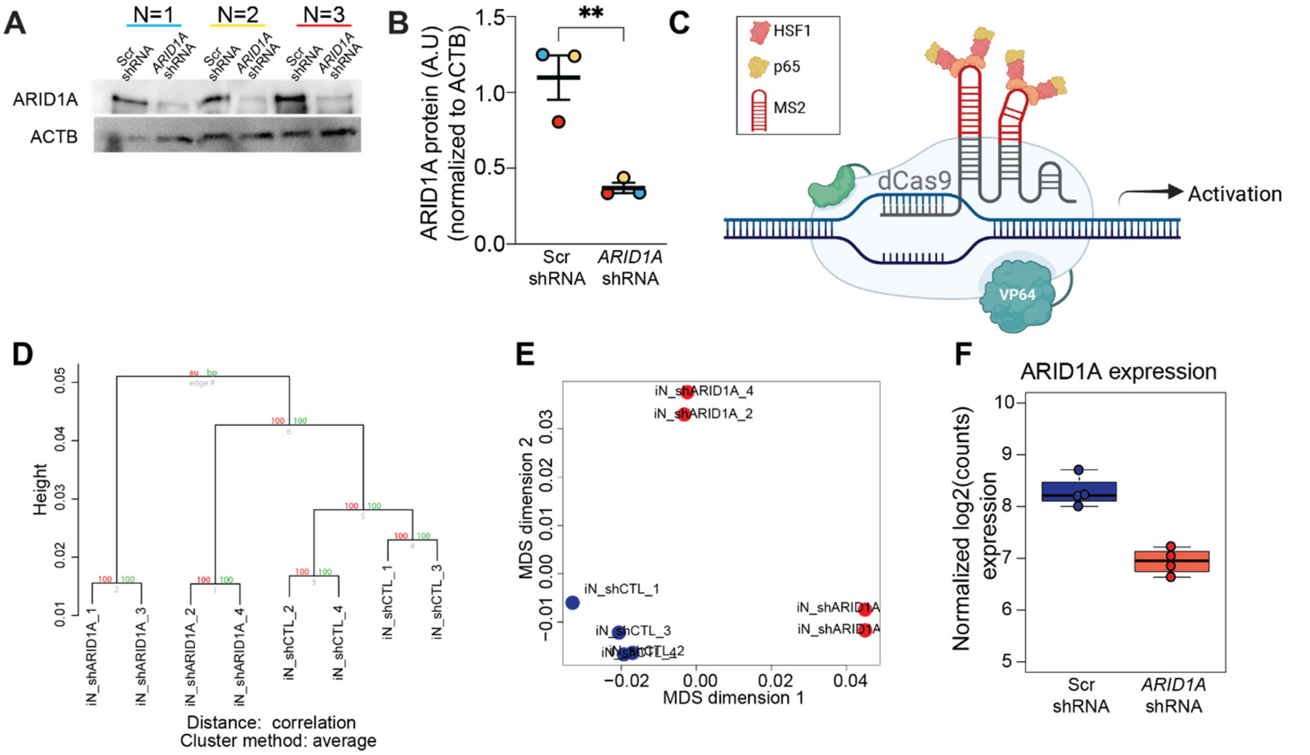


**Fig. S4: *ARID1A* CRISPRa and KD**

A-B) HEK293FT were transduced with either *ARID1A* shRNA or Scr shRNA viral particles for 5 days and collected for protein lysates. ARID1A protein was significantly decreased with *ARID1A* shRNA as compared to Scr ctrl, and normalized to beta actin (ACTB). N=3. Mean ± SEM; Unpaired Student t-test; Arbitrary units (A.U). C) Schematic of CRISPRa experiment (Origene) using dCas9 with enhancers, and targeted gRNAs. D) *ARID1A* shRNA vs Scr shRNA RNA-seq cluster dendrogram reveals increased similarity within group than across groups. E) Multi-dimensional scaling of *ARID1A* shRNA vs Scr shRNA RNA-seq visualizing all replicates. F) *ARID1A* transcript expression in RNA-seq (log2(counts)) is reduced in ARID1A shRNA KD vs. Scr ctrl. *p<0.05 **p<0.01 ***p<0.001 ****p<0.0001.
